# Coupling of death receptor p75^NTR^ to the RhoA and NF-kB pathways differentially regulates internalization in hippocampal and cerebellar neurons

**DOI:** 10.64898/2026.02.10.705215

**Authors:** Xuetong Li, Ziyi Feng, Ajeena Ramanujan, Meng Xie, Carlos F. Ibáñez

## Abstract

Receptor internalization regulates the duration and qualitative output of intracellular signaling. Although generic mechanisms of receptor internalization are well characterized, how these are deployed and regulated in different cell types remains much less understood. Here we show that the p75 neurotrophin receptor (p75^NTR^), a key regulator of neuron survival and function, internalizes in hippocampal (HCNs)and cerebellar granule (CGNs)neurons with very different kinetics, regulated by distinct mechanisms. Compared to HCNs, p75^NTR^ internalizes at a much slower rate and shows stronger interaction with caveolin in CGNs. In both cell types, p75^NTR^ internalization was enhanced by nerve growth factor (NGF)but reduced by inhibitors of the RhoA and PKC pathways. In line with this, internalization of a p75^NTR^ mutant specifically impaired in RhoA signaling was significantly reduced and insensitive to NGF in both neuron types. Accordingly, this mutant showed a much stronger interaction with caveolin than wild type p75^NTR^. On the other hand, internalization of a mutant specifically impaired in coupling to the NF-kB signaling pathway was greatly accelerated in CGNs but unaffected in HCNs. These results reveal the crucial role of intracellular signaling in p75^NTR^ internalization and demonstrate that the receptor is differentially wired to the endocytosis machinery in different neuron types, leading to distinct internalization behaviors.

## Introduction

Cell surface receptors are internalized in vesicles into the cell cytoplasm both spontaneously and upon ligand stimulation in a process known as endocytosis. The majority of spontaneously internalized receptors are normally recycled back to the plasma membrane in vesicles marked by the small GTPases Rab11 or Rab4 (Gruenberg, 2001; Pfeffer, 2003; Roy and Wrana, 2005). Ligand stimulation enhances receptor internalization and alters the intracellular trafficking of endocytic vesicles, also known as endosomes, away from the recycling pathway and deeper into the cell interior on vesicles marked by Rab7 (Deinhardt et al., 2007; Sorkin and Zastrow, 2009). In most cases, such endosomes will eventually fuse with lysosomes for degradation or multivesicular bodies for further sorting to e.g. extracellular vesicles (Bartheld and Altick, 2011). Receptor endocytosis regulates both the duration and quality of receptor signaling as endosomes travel inside the cell and encounter different cellular components (Miaczynska et al., 2004; Sorkin and Zastrow, 2009). Thus, receptor signaling both regulates and is regulated by its own internalization. Endocytosis and intracellular trafficking of cell surface receptors place the biochemistry of receptor signaling in a cellular context. In order to fully understand receptor signaling it is therefore crucial to elucidate how different cellular environments both within the same cell and among different cell types affect receptor internalization and intracellular trafficking. In the central nervous system (CNS), with its great variety of cell types, such level of understanding is currently lacking.

The p75 neurotrophin receptor (p75^NTR^)is expressed by a variety of neurons and glial cells in the peripheral and central nervous systems where it regulates multiple aspects of their development, function and response to injury and degenerative processes, such as Alzheimer’s disease (Ibáñez and Simi, 2012; Roux and Barker, 2002; Underwood and Coulson, 2008). In addition to neurotrophins such as nerve growth factor (NGF)and brain-derived neurotrophic factor (BDNF), p75^NTR^ can also interact with a variety of other ligands, including myelin components such as myelin-associated glycoprotein (MAG), amyloid-beta and viral capsid proteins (Ibáñez and Simi, 2012; Roux and Barker, 2002; Underwood and Coulson, 2008). Three major signaling cascades have been identified downstream of p75^NTR^, namely the RhoA, NF-kB and JNK pathways (Ibáñez and Simi, 2012; Roux and Barker, 2002; Underwood and Coulson, 2008). p75^NTR^ links to RhoA indirectly through its interaction with RhoGDI (Rho GDP-dissociation inhibitor)(Yamashita et al., 1999; Yamashita and Tohyama, 2003), to NF-kB through the binding of RIP2 to the receptor death domain (Carter et al., 1996; Khursigara et al., 2001), and to JNK through several mechanisms, including binding to NRIF or TRAF6 (Friedman, 2000; Khursigara et al., 1999; Casademunt et al., 1999). Other proteins known to interact with the 75^NTR^ intracellular domain include Bex1 (Vilar et al., 2006), SC1 (a.k.a. ARMS)(Chittka and Chao, 1999) and NRAGE (Salehi et al., 2000) among others.

p75^NTR^ internalization and intracellular trafficking has been studied in some detail in peripheral neurons, including sympathetic neurons, as well as cell lines, such as the pheochromocytoma PC12 [reviewd in (Bronfman et al., 2014)]. Studies by Bronfman and colleagues have documented a rather slow p75^NTR^ internalization kinetics (T_1/2_ ≈ 45 min)and a major involvement of the clathrin-dependent endocytic pathway in these cell types (Escudero et al., 2014; Bronfman et al., 2003). A number of studies have also reported on the retrograde transport of p75^NTR^-containing vesicles along the axons of sensory and sympathetic neurons [reviewed in (Ibáñez, 2007; Zweifel et al., 2005)]. Work by the Schiavo laboratory examined p75^NTR^ internalization and retrograde transport in spinal cord motoneurons, reporting spontaneous clathrin-independent recycling but clathrin-dependent retrograde transport in response to NGF (Lalli and Schiavo, 2002; Deinhardt et al., 2007). In comparison, much less is known about these processes in brain neurons. In an earlier study from our laboratory we reported on the internalization kinetics of p75^NTR^ in embryonic mouse hippocampal neurons, and found this to be considerably faster (T_1/2_ ≈ 5min)than that described in peripheral neurons receptor (Yi et al., 2021). Interestingly, receptor mutants deficient in downstream signaling –either lacking the death domain (ΔDD)or with transmembrane Cys^259^ replaced with Ala (C259A)– showed a much slower internalization kinetics in these neurons (T_1/2_ ≈ 12min), associated more readily with Rab11^+^ recycling vesicles, and showed no binding to β-adaptin, a component of clathrin-mediated internalization, compared to the wild type receptor (Yi et al., 2021). These results suggested an important role of p75^NTR^ signaling in controlling its internalization and intracellular trafficking. Importantly, however, which signaling pathways affect p75^NTR^ internalization and whether similar processes and mechanisms apply to p75^NTR^ in neurons from other brain regions is currently unknown.

To fill this knowledge gap, we set out to compare the internalization kinetics of p75^NTR^ in two very different classes of brain neurons, namely hippocampal neurons (herein referred to as HCNs)and cerebellar granule neurons (i.e. CGNs), isolated from late embryonic or early postnatal stages, respectively, of mouse brain development. We discovered that p75^NTR^ is internalized with very different kinetics in these two cell types. Looking for possible mechanistic bases for such difference, we uncovered the distinct roles of the RhoA and NF-kB signaling pathways in the internalization and intracellular trafficking of p75^NTR^, thereby illuminating the crucial role of intracellular signaling in receptor internalization and how this contributes to different internalization behaviors.

## Results

### Distinct p75^NTR^ internalization kinetics in HCNs and CGNs

We compared the internalization kinetics of endogenous p75^NTR^ in HCNs (embryonic day E17.5) and CGNs (postnatal day P3)of wild type C57BL/6 mice. At basal conditions, without ligand stimulation, p75^NTR^ internalized at a much slower rate in CGNs (T_1/2_ ≈ 20min)compared to HCNs (T_1/2_ ≈ 7.8min), while similar maximal levels were reached in both cell types 60min after the onset of internalization (Figure 1A). It has been reported that the faster internalization rate of p75^NTR^ in HCNs compared to sympathetic neurons may be due to the much lower cholesterol content in the former (Bronfman et al., 2014). However, cholesterol levels in HCNs and CGNs were quite comparable in these cultures (Figure S1A-C), indicating that differences in their p75^NTR^ internalization kinetics cannot be accounted for by dissimilar cholesterol contents in these neuron types. As previously shown in peripheral neurons (Bronfman et al., 2003), spontaneous p75^NTR^ internalization was accelerated in the presence of NGF (Figure 1B and C). Despite their very different internalization kinetics at basal conditions, both neuron types responded in a similar fashion to NGF, reducing their T_1/2_ of p75^NTR^ internalization following ligand stimulation (Figure 1B and C). In addition, in both HCNs and CGNs, NGF treatment altered the trafficking of internalized p75^NTR^ away from recycling Rab11^+^ endosomes and into late Rab7^+^ endosomes (Figure 1D and E), in agreement with previous findings in other cell types [e.g. (Deinhardt et al., 2007)].

**Figure 1.**
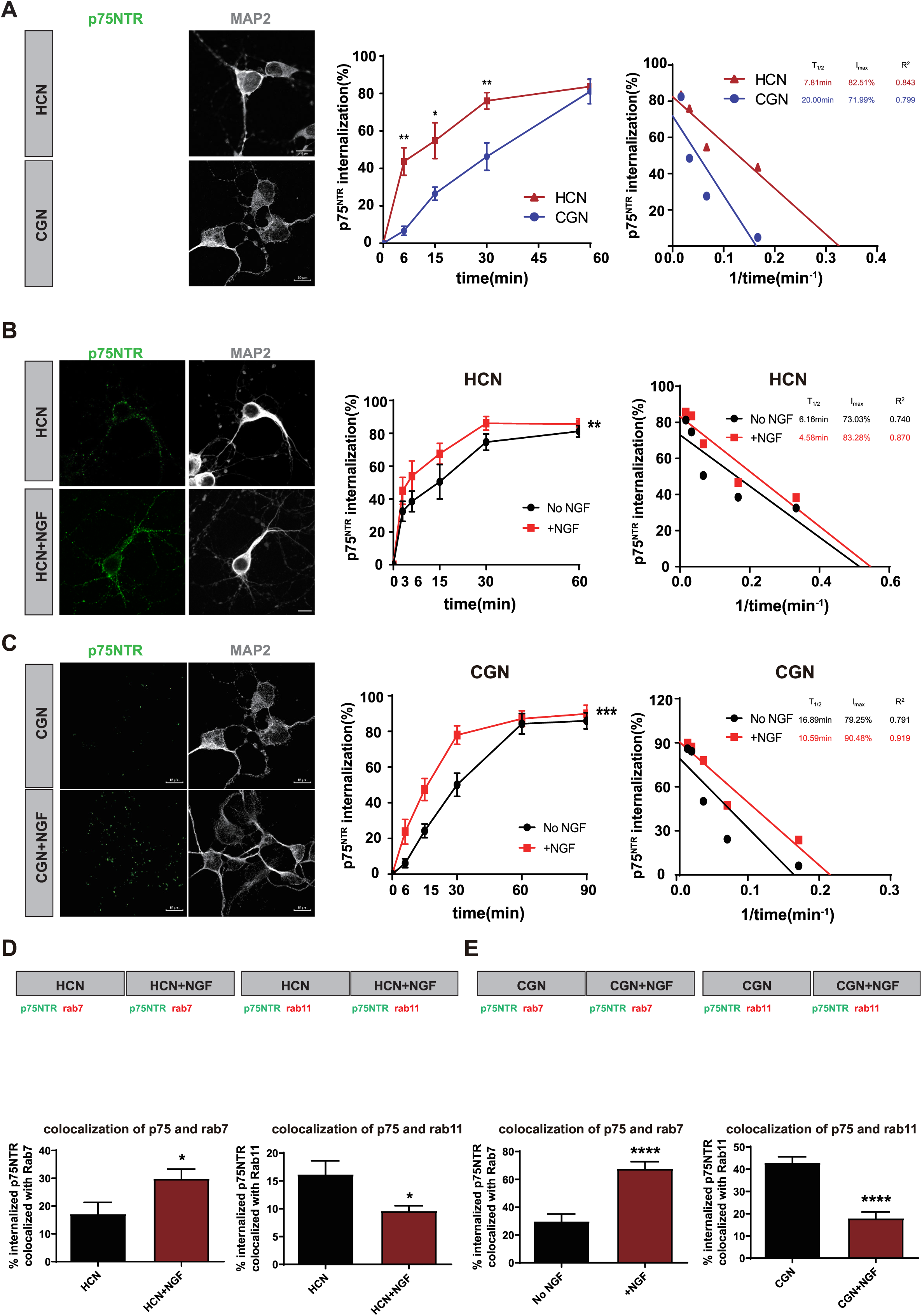
p75^NTR^ shows distinct internalization kinetics in HCNs and CGNs. (A) Distinct p75^NTR^ internalization kinetics in HCNs and CGNs. Representative images show p75^NTR^ after 15min internalization in HCNs and CGNs, respectively. Scale bar, 10μm. Shown is mean±SEM of percentage internalization of total surface p75^NTR^ (set to 100%). N= more than 7 independent experiments. Two-way ANOVA followed by Tukey’s multiple comparisons test. (B) Effects of NGF on p75^NTR^ internalization in HCNs. Representative images show p75^NTR^ after 6min internalization in the presence or absence of 50ng/mL NGF. Scale bar, 10μm. Shown is mean±SEM of percentage internalization of total surface p75^NTR^ (set to 100%). N= more than 5 independent experiments. Two-way ANOVA followed by Tukey’s multiple comparisons test. (C) Effects of NGF on p75^NTR^ internalization in CGNs. Representative images show p75^NTR^ after 15min internalization in the presence or absence of 50ng/mLNGF. Scale bar, 10μm. Shown is mean±SEM of percentage internalization of total surface p75^NTR^ (set to 100%). N= more than 7 independent experiments. Two-way ANOVA followed by Tukey’s multiple comparisons test. (D) Effects of NGF on p75^NTR^ intracellular trafficking in HCNs. Representative images show colocalization of internalized p75^NTR^ and Rab7 or Rab11 after 30min internalization in HCNs. Scale bar, 10μm. Histograms show the percentage of internalized p75^NTR^ colocalized with rab7 or rab11 as indicated. Shown is mean±SEM. N=3 independent experiments. Unpaired t test with Welch’s correction. (E) Effects of NGF on p75^NTR^ intracellular trafficking in CGNs. Effects of NGF on p75^NTR^ intracellular trafficking in CGNs. Representative images show colocalization of internalized p75^NTR^ and Rab7 or Rab11 after 60min internalization in CGNs. Scale bar, 10μm. Histograms show the percentage of internalized p75^NTR^ colocalized with rab7 or rab11 as indicated. Shown is mean±SEM. N=3 independent experiments. Unpaired t test with Welch’s correction.

### Contrasting interaction of internalized p75^NTR^ with clathrin- and caveolin-containing vesicles in HCNs vs. CGNs

Using the Protein Ligation Assay (PLA), we assessed the interaction between internalized p75^NTR^ and components of the clathrin and caveolin endocytosis machineries in HCNs and CGNs. We found that internalized p75^NTR^ interacted with AP2B1 (a subunit of the clathrin-associated adaptor protein complex)to a lower level in CGNs than in HCNs (Figure 2A). In contrast, the receptor interacted at a higher level with caveolin-1 in CGNs than in HCNs after internalization (Figure 2B). This result is in agreement with the slower internalization of p75^NTR^ in CGNs, as it has been proposed that caveolin-mediated internalization of cell surface receptors is slower than clathrin-mediated internalization (Mazumdar et al., 2021). In both neuron types, NGF treatment enhanced internalized p75^NTR^ binding to AP2B1 (Figure 2C and D), but diminished the interaction of internalized p75^NTR^ with caveolin-1 in CGNs, while a small but below statistical significance decrease was also observed in HCNs (Figure 2E and F). This result is also in agreement with the effects of NGF on the speed of p75^NTR^ internalization. In both HCNs and CGNs, p75^NTR^ internalization was completely abolished after inhibition of dynamin, the essential mediator of vesicle excision from the plasma membrane in both clathrin- and caveolin-mediated endocytosis (Figure 2G). Together, these data indicate a differential interaction of internalized p75^NTR^ with clathrin-vs. caveolin-containing vesicles in HCNs and CGNs, but comparable effects of ligand treatment on the interaction of the internalized receptor with the endocytosis machinery in the two neuron types.

**Figure 2.**
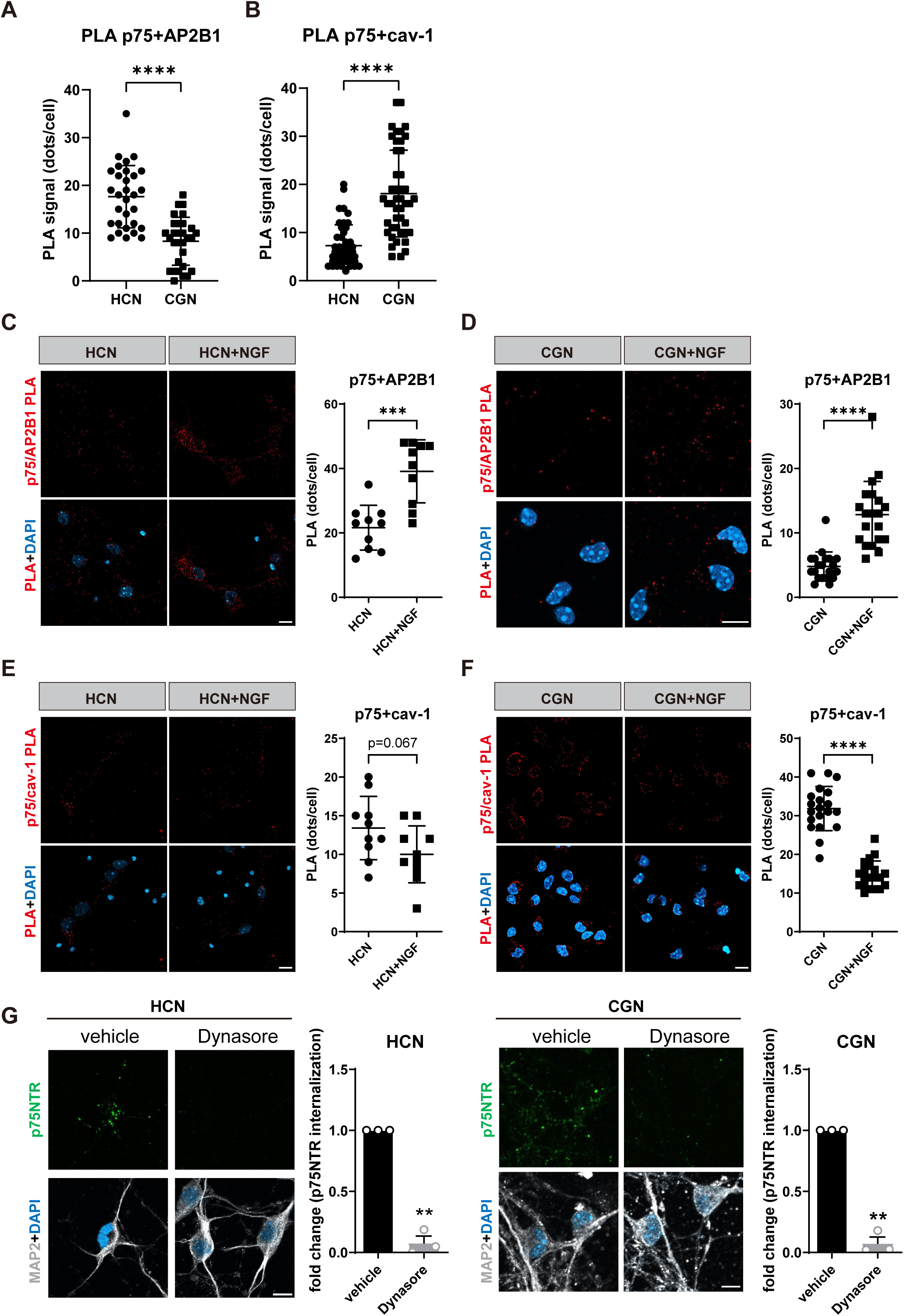
Contrasting p75^NTR^ interactions with caveolin- and clathrin-containing vesicles in HCNs and CGNs. (A) Quantification of PLA signals between p75^NTR^ and AP2B1 in HCNs and CGNs, as indicated. Values were shown as mean±SD PLA puncta per cell from 30 neurons in 3 independent experiments. One-way ANOVA followed by Tukey’s multiple comparisons test. (B) Quantification of PLA signals between p75^NTR^ and caveolin-1 in HCNs and CGNs, as indicated. Values were shown as mean±SD PLA puncta per cell from 45 neurons in 3 independent experiments. Unpaired t test with Welch’s correction. Unpaired t test with Welch’s correction. (C) PLA assay between p75^NTR^ and AP2B1 in HCNs in the presence or absence of NGF, as indicated. Scale bar, 10μm. Values were shown as mean±SD PLA puncta per cell from 10 neurons from duplicate coverslips. Unpaired t test with Welch’s correction. (D) PLA assay between p75^NTR^ and AP2B1 in CGNs in the presence or absence of NGF, as indicated. Scale bar, 10μm. Values were shown as mean±SD PLA puncta per cell from 20 neurons in 2 independent experiments. Unpaired t test with Welch’s correction. (E) PLA assay between p75^NTR^ and caveolin-1 in HCNs in the presence or absence of NGF, as indicated. Scale bar, 10μm. Values were shown as mean±SD PLA puncta per cell from 10 neurons from duplicate coverslips. Unpaired t test with Welch’s correction. (F) PLA assay between p75^NTR^ and caveolin-1 in CGNs in the presence or absence of NGF, as indicated. Scale bar, 10μm. Values were shown as mean±SD PLA puncta per cell from 20 neurons in 2 independent experiments. Unpaired t test with Welch’s correction. (G) Effects of Dynasore on p75^NTR^ internalization after 60min in HCNs and CGNs. Scale bar, 10μm. Shown is mean±SEM of p75^NTR^ internalization normalized to the average of vehicle group. N= 3 independent experiments. Unpaired t test with Welch’s correction.

### Reduced p75^NTR^ internalization in HCNs and CGNs after pharmacological inhibition of the RhoA pathway

Our previous studies showed that signaling-deficient p75^NTR^ mutants ΔDD and C259A display slower and reduced internalization in HCNs compared to the wild type receptor (Yi et al., 2021), highlighting the importance of downstream signaling in the internalization mechanism of p75^NTR^. In order to gain insights into the specific signaling pathways that regulate p75^NTR^ internalization in HCNs and CGNs, we investigated the effects of a battery of pharmacological inhibitors on the internalization behavior of the receptor in the two neuron types. We found that neither the JNK inhibitor JNK-IN-8, nor the pan-caspase inhibitor Q-VD-OpH, nor JSH-23, an inhibitor of the nuclear translocation of the p65^NF-kB^ subunit, had significant effects on p75^NTR^ internalization in either neuron type (Figure 3A and B). On the other hand, both the RhoA inhibitor Rhosin and ROCK inhibitor Fasudil diminished the internalization of p75^NTR^ in both HCNs and CGNs (Figure 3A and B), indicating the requirement of a functional RhoA pathway in the endocytosis of this receptor. p75^NTR^ internalization was also greatly diminished upon inhibition of Protein Kinase C (PKC) with Gö6976 (Figure 3C). This result further supports the involvement of RhoA signaling in p75^NTR^ internalization, as our previous work demonstrated that phosphorylation of RhoGDI at Ser^34^ by PKC greatly enhances RhoGDI binding to the receptor, leading to increased RhoA activation (Ramanujan and Ibáñez, 2024).

**Figure 3.**
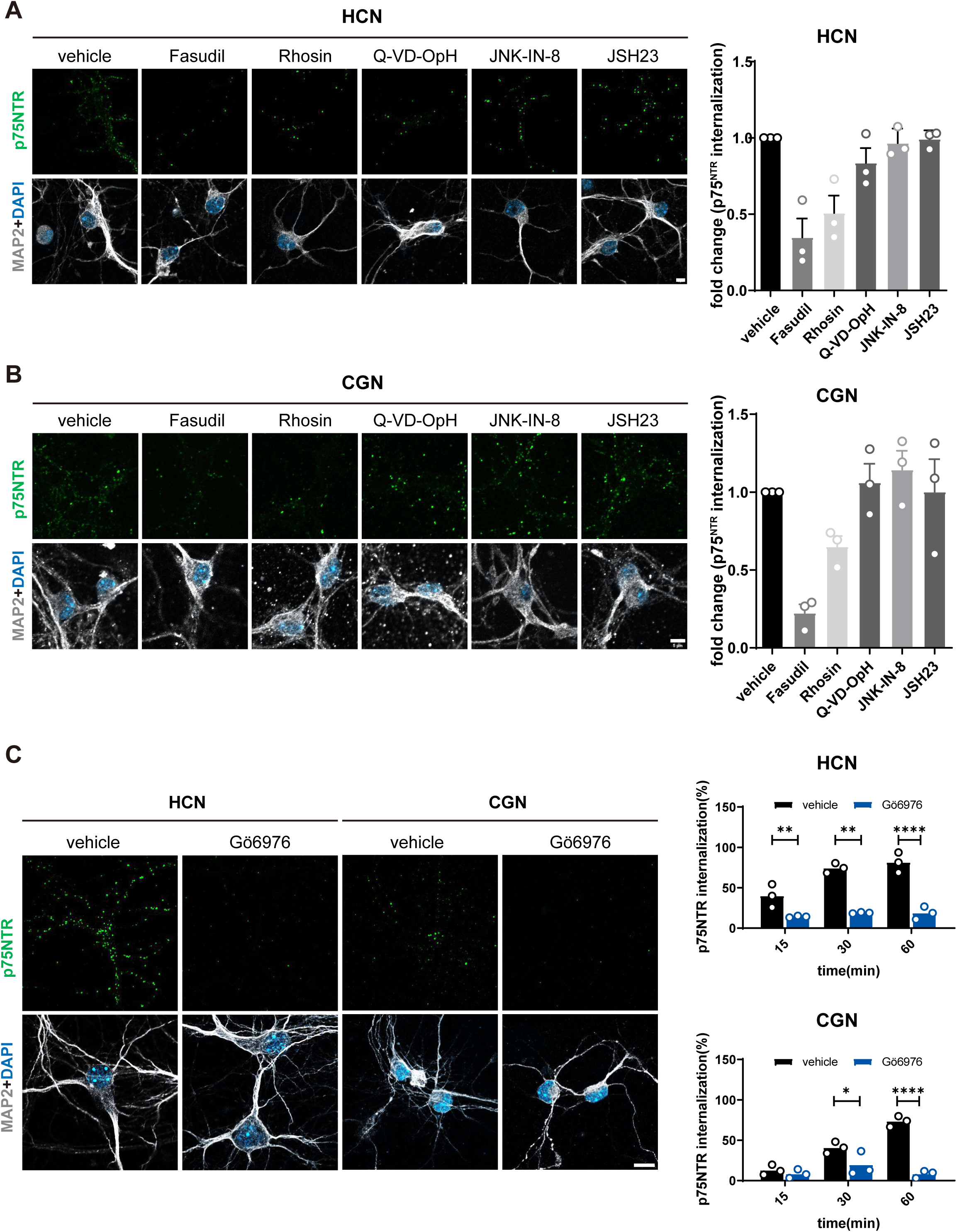
Reduced p75^NTR^ internalization in HCNs and CGNs after pharmacological inhibition of the RhoA pathway, but not JNK or NF-kB pathways. (A) Effects of signaling inhibitors on p75^NTR^ internalization after 60min in HCNs. Scale bar, 5μm. Shown is mean±SEM of p75^NTR^ internalization normalized to the average of vehicle group. N= 3 independent experiments. One-way ANOVA followed by Tukey’s multiple comparisons test. (B) Effects of signaling inhibitors on p75^NTR^ internalization after 60min in CGN. Scale bar, 5μm. Shown is mean±SEM of p75^NTR^ internalization normalized to the average of vehicle group. N= 3 independent experiments. One-way ANOVA followed by Tukey’s multiple comparisons test. (C) Effects of PKC inhibitor Gö6976 on p75^NTR^ internalization in HCNs and CGNs. Representative images show p75^NTR^ after 60min internalization in the presence or absence of Gö6976. Scale bar, 10μm. Shown is mean±SEM of percentage internalization of total surface p75NTR (set to 100%). N= 3 independent experiments. Two-way ANOVA followed by Tukey’s multiple comparisons test.

### A p75^NTR^ mutant that is specifically uncoupled from the RhoGDI/RhoA pathway

Based on the effects of RhoA, ROCK and PKC inhibitors on the internalization of p75^NTR^, we sought to specifically uncouple the receptor from this pathway. To this end, we looked for intracellular domain mutants that would specifically abolish the binding of RhoGDI to the receptor, while retaining interactions with mediators of other signaling pathways. In previous work, we identified Lys^303^ (mouse p75^NTR^ numbering) in the juxtamembrane domain of p75^NTR^ intracellular region as a critical determinant for the interaction of the receptor with phosphorylated Ser^34^ in RhoGDI, following phosphorylation of the latter by PKC (Ramanujan and Ibáñez, 2024). Although binding of p75^NTR^ to RhoGDI was greatly reduced in a mutant receptor with Alanine replacing Lys^303^ (K303A) (Ramanujan and Ibáñez, 2024), the interaction was not completely abolished. As RhoGDI also interacts with residues in the p75^NTR^ death domain (DD), we sought to identify additional mutations in this domain that would completely but specifically abolished this interaction. In earlier work, we performed an alanine-scanning structure-function analysis of the p75^NTR^ DD and identified several solvent-accessible residues that mediate interaction of the receptor with components of different signaling pathways, including the RhoGDI/RhoA, RIP2/NF-kB and JNK/cell death pathways (Charalampopoulos et al., 2012). However, while we could identify mutants that specifically uncoupled the receptor from the RIP2/NF-kB and JNK/cell death pathways, the mutants that abolished DD binding to RhoGDI also affected one of the other two pathways (Charalampopoulos et al., 2012). With the subsequent determination of the 3D NMR structure of the complex between the p75^NTR^ DD and RhoGDI by our group (Lin et al., 2015), it became possible to undertake a more specific mutagenesis analysis of DD residues directly involved in its interaction with RhoGDI. Guided by this NMR structure, we identified DD residues Lys^346^ and Glu^349^ as important contributors to p75^NTR^ binding to RhoGDI. Co-immunoprecipitation experiments in transfected NIH3T3 cells showed that Alanine mutation of either residue (K346A and E349A, respectively), individually or in conjunction with the K303A mutation, greatly diminished the interaction between p75^NTR^ and RhoGDI (Figure 4A). Importantly, a triple receptor mutant with all three residues changed to Alanine (K303A/K346A/E349A) showed no detectable binding to RhoGDI (Figure 4A). All these mutants showed comparable levels of expression in transfected cells (Figure S2). The functional significance of these results was confirmed by assessing the relative contributions of these residues to the activation of RhoA in NIH3T3 cells expressing the different p75^NTR^ mutants (Figure 4B). As reported previously (Ramanujan and Ibáñez, 2024), overexpression of wild type p75^NTR^ greatly enhances activation of endogenous RhoA in these cells, as assessed by the levels of RhoA bound to GTP (Figure 4B). While individually or in pairs, the p75^NTR^ mutants elicited lower RhoA activation, the triple mutant (herein termed p75^KKEA^ or KKEA) showed no detectable stimulation of RhoA activity over background in this assay (Figure 4B).

**Figure 4.**
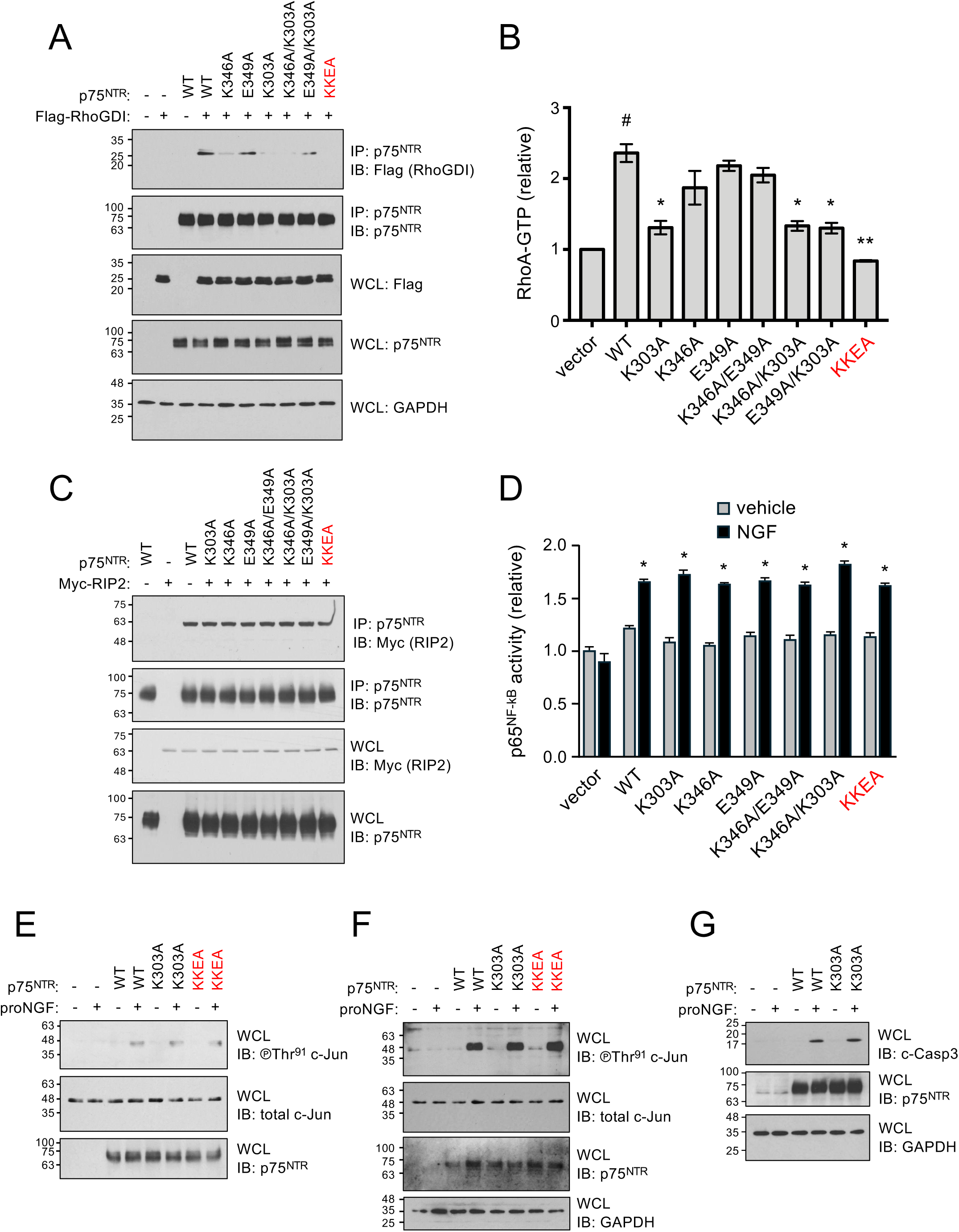
A mutant of p75^NTR^ that is specifically uncoupled from the RhoGDI/RhoA pathway. (A) Triple alanine replacement of Lys^303^, Lys^346^ and Glu^349^ (KKEA)in the p75^NTR^ intracellular domain completely abolishes binding to RhoGDI. Co-immunoprecipitation analysis of interaction between wild type and mutant p75^NTR^ (as indicated)with Flag-tagged RhoGDI in transfected NIH3T3 cells (top panel). The lower panels are controls for immunoprecipitation and loading. IP, immunoprecipitation; IB: immunoblotting; WCL: whole cell lysate. (B) KKEA triple p75^NTR^ mutant (p75^KKEA^)fails to enhance RhoA GTPase activity. RhoA-GTP assay (G-LISA)in NIH3T3 cells transfected with the indicated constructs. Mean ± SEM of reading at 490 nm from 3 independent experiments are shown. #, P<0.0001 vs. vector; **, P<0.01 and *, P<0.05 vs. WT; one-way ANOVA followed by Tukey’s multiple comparison test. (C) KKEA mutation does not affect binding of p75^NTR^ to RIP2. immunoprecipitation analysis of interaction between wild type and mutant p75^NTR^ (as indicated)with Myc-tagged RIP2 in transfected NIH3T3 cells (top panel). The lower panels are controls for immunoprecipitation and loading. IP, immunoprecipitation; IB: immunoblotting; WCL: whole cell lysate. (D) KKEA mutation does not affect p75^NTR^–mediated activation of p65^NF-kB^ in response to 50ng/mL NGF. CGNs lacking p75^NTR^ (knock-out)were reconstituted with the indicated constructs via lentivirus transduction. Nuclear p65-NFkB localization is measured spectrophotometrically at 450nm (see Methods). Results are shown as mean ± SEM of triplicate readings normalized to vector (vehicle). *, P<0.05 vs. corresponding vehicle; one-way ANOVA followed by Tukey’s multiple comparison test. (E) KKEA mutation does not affect p75^NTR^–mediated activation of c-Jun in response to proNGF in NIH3T3 cells. Western blot analysis of activated c-Jun (phosphorylated at Thr^91^)after transfection of the indicated p75^NTR^ constructs in the presence and absence of 2.5ng/mL proNGF (top panel). The lower panels are controls for loading. IB: immunoblotting; WCL: whole cell lysate. (F) KKEA mutation does not affect p75^NTR^–mediated activation of c-Jun in response to proNGF in CGNs. Western blot analysis of activated c-Jun (phosphorylated at Thr^91^)after lentiviral transduction of the indicated p75^NTR^ constructs into p75^NTR^ knockout CGNs in the presence and absence of 2.5ng/mL proNGF (top panel). The lower panels are controls for loading. IB: immunoblotting; WCL: whole cell lysate. (G) KKEA mutation does not affect p75^NTR^–mediated activation of caspase-3 in response to proNGF in CGNs. Western blot analysis of activated caspase-3 (cleaved caspase-3)after lentiviral transduction of the indicated p75^NTR^ constructs into p75^NTR^ knockout CGNs in the presence and absence of 2.5ng/mL proNGF (top panel). The lower panels are controls for loading. IB: immunoblotting; WCL: whole cell lysate.

In order to determine the specificity of the effects of these mutants on the RhoA pathway, we examined their ability to affect the RIP2/NF-kB and JNK/cell death pathways in comparison to the wild type receptor. Co-immunoprecipitation experiments in transfected NIH3T3 cells showed that the different p75^NTR^ mutants that affected RhoGDI binding and RhoA activation, including triple mutant KKEA, had no effect in the interaction of the receptor with RIP2, a key downstream effector linking p75^NTR^ to NF-kB signaling (Figure 4C). Next, we assessed activation of NF-kB in CGNs lacking p75^NTR^ (KO CGNs) after lentiviral-mediated transduction of wild type and mutant p75^NTR^ molecules. While transduction of wild type p75^NTR^ did not per se affected NF-kB activation, this was greatly stimulated upon NGF treatment (Figure 4D). NGF had no effects in KO CGNs (Figure 4D). Importantly, the levels of NF-kB activation elicited by NGF stimulation of p75^NTR^ mutants remained similar to that of the wild type receptor (Figure 4D), indicating that none of the mutations, including KKEA, affected the ability of p75^NTR^ to regulate the activation of NF-kB. Finally, we examined activation of the JNK/cell death pathway by assessing phosphorylation of c-Jun, a direct target of JNK, and caspase-3 cleavage, a key determinant of apoptotic cell death. In NIH3T3 cells, treatment with proNGF (which as opposed to NGF is a potent stimulator of the JNK/cell death pathway)induced phosphorylation of c-Jun at Thr^91^ in cells transfected with wild type p75^NTR^ plasmid but not with empty vector (Figure 4E). Importantly, comparable levels of c-Jun phosphorylation were observed in cells that received plasmids encoding the K303A or KKEA mutants (Figure 4E). Similar results were obtained in KO CGNs that were reconstituted with wild type or mutant p75^NTR^ receptors by lentiviral transduction (Figure 4F). Likewise, comparable levels of cleaved caspase-3 were observed in KO CGNs expressing wild type or mutant p75^NTR^ following treatment with proNGF (Figure 4G). Together, these results indicate that the KKEA mutation specifically affects the ability of p75^NTR^ to couple to the RhoA/RhoGDI pathway without affecting the RIP2/NF-kB or JNK/cell death pathways.

We generated knock-in mice carrying the KKEA mutation in the *Ngfr* locus, encoding p75^NTR^, and verified that the mutant receptor deficient in RhoA signaling was expressed at levels comparable to wild type (Figure S3A and 3B). Using the PLA assay, we assessed the interaction of p75^NTR^ with RhoGDI in HCNs and CGNs isolated from wild type and KKEA mutant mice. Binding of p75^KKEA^ to RhoGDI was dramatically reduced in mutant HCNs and CGNs (Figure 5A and B). In the latter, PMA (a potent stimulator of PKC activity)increased the interaction of p75^NTR^ with RhoGDI in wild type but not in KKEA mutant CGNs (Figure 5B). In HCNs, no PLA signal between WT p75^NTR^ and RIP2 can be observed due to absent (or very low) RIP2 expression (Figure S3C), in agreement with our earlier study (Vicario et al., 2015). With regards to the cell death pathway, CGNs from p75^KKEA^ mice were as sensitive to proNGF-induced elevation in the levels of activated caspase-3 as neurons expressing WT p75^NTR^ (Figure 5D), which confirms our observations in transfected NIH3T3 cells and KO neurons. Together, our data indicate that knock-in p75^KKEA^ mutant mice carry a mutant p75^NTR^ receptor that is unable to couple to the RhoGDI/RhoA pathway.

**Figure 5.**
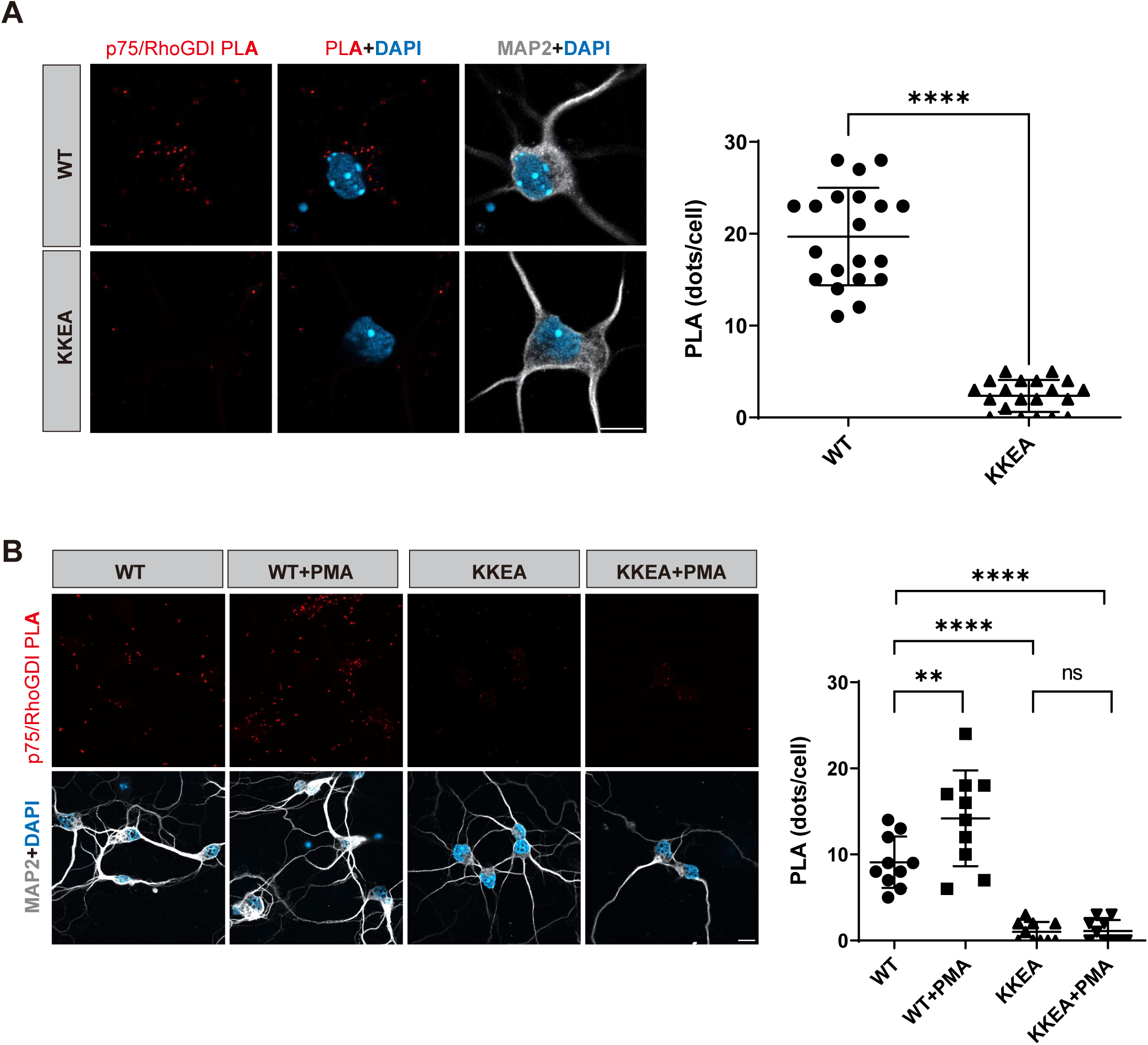
Lack of binding of p75^NTR^ to RhoGDI in HCNs and CGNs from p75^KKEA^ mutant mice. (A) Abolished binding of p75^NTR^ to RhoGDI in HCNs of p75^KKEA^ mutant mice. PLA assay between p75^NTR^ and RhoGDI in HCNs. Scale bar, 10μm. Values were shown as mean±SD PLA puncta per cell from 20 neurons from duplicate coverslips. Unpaired t test with Welch’s correction. (B) Abolished binding of p75^NTR^ to RhoGDI in CGNs of p75^KKEA^ mutant mice. PLA assay between p75^NTR^ and RhoGDI in CGNs. PMA was added to either WT or KKEA CGNs at xμM. Scale bar, 10μm. Values were shown as mean±SD PLA puncta per cell from 10 neurons from duplicate coverslips. Two-way ANOVA followed by Tukey’s multiple comparisons test

### Impaired p75^NTR^ internalization in HCNs and CGNs from p75^KKEA^ mutant mice

Analyses of p75^NTR^ internalization in HCNs and CGNs isolated from p75^KKEA^ mutant mice showed a severe impairment in both neuron types, both in the speed as well as the maximal levels reached (Figure 6A and B), indicating that coupling to RhoGDI/RhoA signaling is crucial for p75^NTR^ internalization. Interestingly, while NGF treatment accelerated internalization of the wild type receptor in both neuron types, it had no effect on p75^KKEA^ mutant neurons (Figure 6A and B). The impairment in the internalization of the p75^KKEA^ receptor was similar to that previously observed (Yi et al., 2021) in signaling-deficient p75^NTR^ mutants lacking the death domain (ΔDD)or with transmembrane Cys^259^ replaced to Ala (C259A) (Figure 6C), suggesting that coupling to RhoA signaling may be the most critical pathway controlling efficient p75^NTR^ internalization.

**Figure 6.**
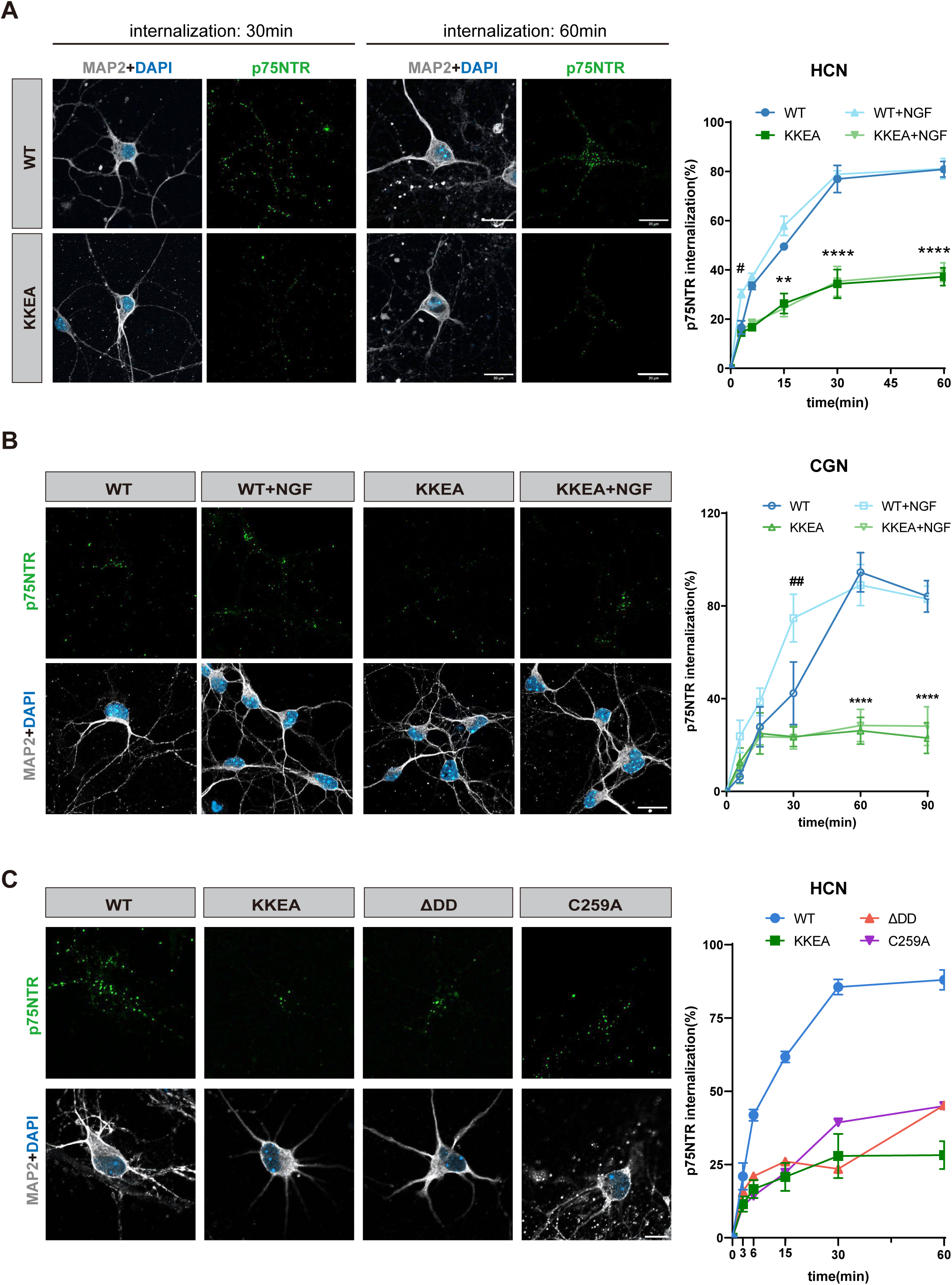
Impaired p75^NTR^ internalization in HCNs and CGNs from p75^KKEA^ mutant mice. (A) p75^NTR^ internalization kinetics in WT and KKEA HCNs in the presence and absence of 50ng/mL NGF. Scale bar, 20μm. Shown is mean±SEM of percentage internalization of total surface p75^NTR^ (set to 100%). N= 3 independent experiments. Two-way ANOVA followed by Tukey’s multiple comparisons test. (B) p75^NTR^ internalization kinetics in WT and KKEA CGNs in the presence and absence of 50ng/mL NGF. Scale bar, 10μm. Shown is mean±SEM of percentage internalization of total surface p75NTR (set to 100%). Representative images show p75^NTR^ after 60min internalization. N= 3 independent experiments. Two-way ANOVA followed by Tukey’s multiple comparisons test. (C) p75^NTR^ internalization kinetics in HCNs from wild type (WT), KKEA, ΔDD and C259A embryos as indicated. Representative images show p75^NTR^ after 60min internalization. Scale bar, 10μm. Shown is mean±SEM of percentage internalization of total surface p75^NTR^ (set to 100%). N= 2-3 independent experiments.

In HCNs, p75^KKEA^ interacted with AP2B1 at levels comparable to wild type (Figure 7A) but showed significantly higher interaction with caveolin-1 (Figure 7B), which is in line with the slower rate and reduced overall internalization of the mutant receptor. In contrast to wild type p75^NTR^, interaction of the p75^KKEA^ mutant with either AP2B1 or caveolin-1 was unaffected by NGF treatment (Supplementary Figure 4A and B), in agreement with the inability of NGF to affect p75^KKEA^ internalization (Figure 6A and B). In CGNs, a small but below statistical significance decrease in the interaction of p75^KKEA^ with AP2B1 was observed compared to wild type (Figure 7C), while the interaction of the mutant with caveolin-1 was much higher than wild type (Figure 7D), a result that again aligns with the its reduced internalization. Similar to HCNs, p75^KKEA^ interaction with either AP2B1 or caveolin-1 was unaffected by NGF treatment (Supplementary Figure 4C and D). Together, these results indicate that uncoupling p75^NTR^ from the RhoGDI/RhoA pathway enhances the interaction of the receptor with the caveolin endocytic machinery and renders receptor internalization refractory to the effects of NGF.

**Figure 7.**
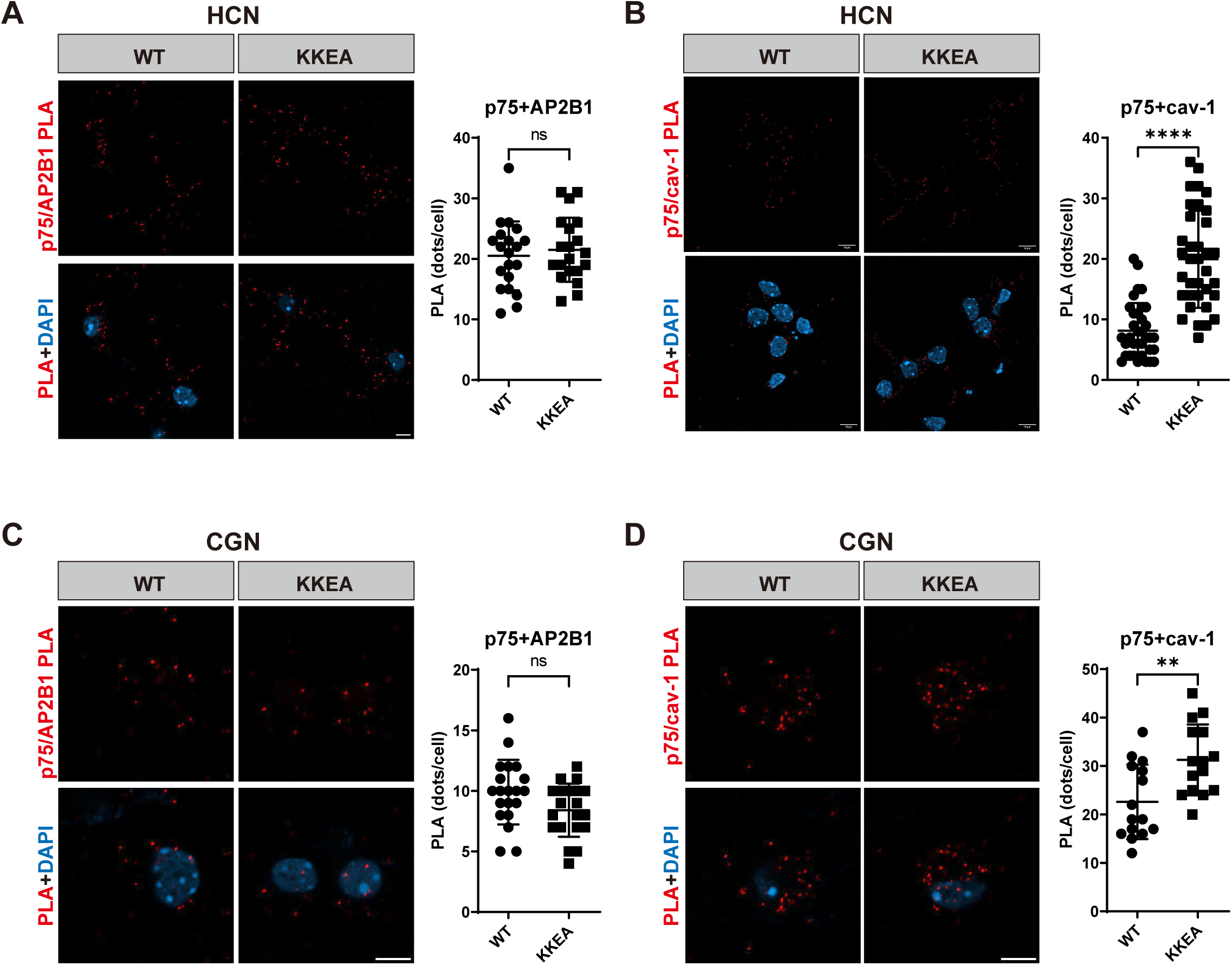
Differential interaction of p75^KKEA^ with components of the clathrin and caveolin endocytosis pathways. (A) PLA assay between p75^NTR^ and AP2B1 in WT and KKEA HCNs. Scale bar, 5μm. Values were shown as mean±SD PLA puncta per cell from 20 neurons in 2 independent experiments. Unpaired t test with Welch’s correction. (B) PLA assay between p75^NTR^ and caveolin-1 in WT and KKEA HCNs. Scale bar, 5μm. Values were shown as mean±SD PLA puncta per cell from 35 neurons in 3 independent experiments. Unpaired t test with Welch’s correction. (C) PLA assay between p75^NTR^ and AP2B1 in WT and KKEA CGNs. Scale bar, 5μm. Values were shown as mean±SD PLA puncta per cell from 20 neurons in 2 independent experiments. Unpaired t test with Welch’s correction. (D) PLA assay between p75^NTR^ and caveolin-1 in WT and KKEA CGNs. Scale bar, 5μm. Values were shown as mean±SD PLA puncta per cell from 15 neurons in 2 independent experiments. Unpaired t test with Welch’s correction.

### Uncoupling p75^NTR^ from the NF-kB pathway accelerates internalization in CGNs but has no effect in HCNs

In addition to RhoA signaling, p75^NTR^ has also the ability to instigate the NF-kB pathway through recruitment of RIP2 to its DD (Lin et al., 2015; Carter et al., 1996; Khursigara et al., 2001). Interestingly, we and others have previously shown that NGF fails to recruit RIP2 to p75^NTR^, stimulate IkB degradation or induce nuclear translocation of p65 NF-kB subunit in HCNs, although it can readily do so in CGNs (Volosin et al., 2008; Vicario et al., 2015), suggesting crucial cell-specific differences in the ability of p75^NTR^ to engage this pathway. This prompted us to investigate whether the ability of p75^NTR^ to link to RIP2 and NF-kB signaling has any impact on its internalization.

In previous work, we characterized three alanine replacements in residues of the p75^NTR^ DD (i.e. Asp^357^, His^361^ and Glu^365^, mouse p75^NTR^ numbering)which together prevented recruitment of RIP2 and upregulation of NF-kB signaling after neurotrophin stimulation, while still being able to couple to RhoA signaling and the JNK/cell death pathway. To uncover physiological roles of this pathway in p75^NTR^ signaling, we generated knock-in mice carrying this triple mutation, herein termed p75^DHEA^ (Supplementary Figure S5A). p75^DHEA^ was expressed at levels comparable to wild type in the postnatal brain of these mice (Supplementary Figure S5B). Since no binding of RIP2 to WT p75^NTR^ can be detected in HCNs (Vicario et al., 2015), we used the PLA assay to only assess the interaction of p75^NTR^ RhoGDI in neurons isolated from wild type and DHEA mutant mice. In these neurons, interaction between p75^DHEA^ and RhoGDI was comparable to that of WT p75^NTR^ and equally sensitive to stimulation with PMA (Figure 8A). On the other hand, binding of p75^DHEA^ to RIP2 was dramatically reduced in CGNs and was unresponsive to NGF (Figure 8B). Interestingly, interaction between p75^DHEA^ and RhoGDI was greatly increased in these neurons compared to WT p75^NTR^ and further enhanced by PMA treatment (Figure 8C). In line with this, the levels of activated RhoA (i.e. RhoA•GTP) were enhanced in DHEA mutant CGNs compared to WT CGNs (Figure 8D). These results were not entirely unexpected. As we have reported elsewhere (Lin et al., 2015), RIP2 and RhoGDI compete for binding to the p75^NTR^ intracellular domain. RIP2 displays much higher affinity than RhoGDI and enhanced RIP2 interaction displaces RhoGDI from the receptor and reduces RhoA activity. Conversely, brain extracts from RIP2 knock-out mice show elevated levels of RhoA•GTP compared to extracts from WT littermates (Lin et al., 2015). Thus the increased interaction of the p75^DHEA^ with RhoGDI is in agreement with our previous findings. With regards to the cell death pathway, CGNs from p75^DHEA^ mice were as sensitive to proNGF-induced elevation in the levels of activated caspase-3 as neurons expressing WT p75^NTR^ (Figure 8E), in agreement with our previous results (Charalampopoulos et al., 2012). We conclude from these studies that p75^DHEA^ mutant mice carry a mutant p75^NTR^ receptor that is unable to couple to the RIP2/NF-kB pathway and, as a result, shows enhanced coupling to the RhoGDI/RhoA pathway.

**Figure 8.**
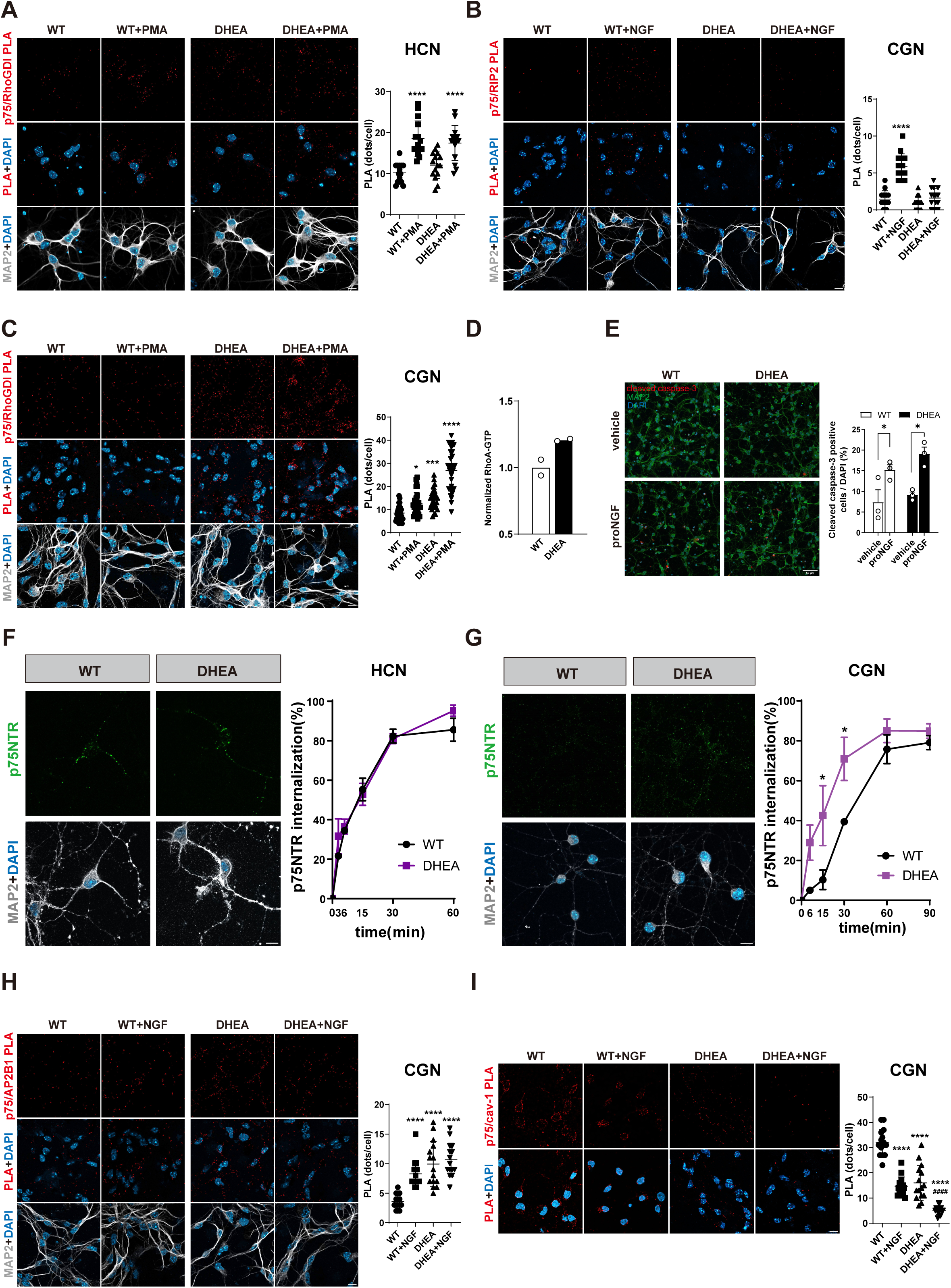
Uncoupling p75^NTR^ from the NF-kB pathway accelerates internalization in CGNs but not in HCNs. (A) Unchanged binding of p75^NTR^ to RhoGDI in HCNs of p75^DHEA^ mutant mice. PLA assay between p75^NTR^ and RhoGDI in HCNs in the presence and absence of PMA (2μM). Scale bar, 10μm. Values were shown as mean±SD PLA puncta per cell from from 3 independent experiments (30 neurons each). Unpaired t test with Welch’s correction. (B) Abolished binding of p75^NTR^ to RIP2 in CGNs of p75^DHEA^ mutant mice. PLA assay between p75^NTR^ and RIP2 in mutant CGNs in the presence and absence of NGF (50ng/ml). Scale bar, 10μm. Values were shown as mean±SD PLA puncta per cell from 3 independent experiments (30 neurons each). Unpaired t test with Welch’s correction. (C) Enhanced binding of p75^NTR^ to RhoGDI in CGNs of p75^DHEA^ mutant mice. PLA assay between p75^NTR^ and RhoGDI in HCNs in the presence and absence of PMA (XμM). Scale bar, 10μm. Values were shown as mean±SD PLA puncta per cell from 3 independent experiments (30 neurons each). Unpaired t test with Welch’s correction. (D) Enhanced RhoA GTPase activity in CGNs of p75^DHEA^ mutant mice. RhoA-GTP assay (G-LISA)in WT and DHEA mutant CGNs. Mean ± SEM of reading at 490 nm from 2 independent experiments are shown. Unpaired t test with Welch’s correction. (E) Unchanged caspase-3 activation by proNGF in CGNs from p75^DHEA^ mutant mice. Shown is activated caspase-3 immunostaining in the presence or absence of proNGF (2.5ng/ml). Quantification shows positive cells normalized to DAPI nuclei as mena±SEM. N=3 independent experiments. Two-way ANOVA followed by Tukey’s multiple comparisons test. (F) p75^NTR^ internalization kinetics in WT and DHEA HCNs. Scale bar, 10μm. Shown is mean±SEM of percentage internalization of total surface p75^NTR^ (set to 100%). N=3 independent experiments. Two-way ANOVA followed by Tukey’s multiple comparisons test. (G) p75^NTR^ internalization kinetics in WT and DHEA CGNs. Scale bar, 10μm. Shown is mean±SEM of percentage internalization of total surface p75^NTR^ (set to 100%). Representative images show p75NTR after 60min internalization. N= 3 independent experiments. Two-way ANOVA followed by Tukey’s multiple comparisons test. (H) PLA assay between p75^NTR^ and AP2B1 in CGNs from WT and p75^DHEA^ mice in the presence or absence of NGF, as indicated. Scale bar, 10μm. Values were shown as mean±SD PLA puncta per cell. N=3 experiments (30 neurons each). Unpaired t test with Welch’s correction. (I) PLA assay between p75^NTR^ and caveolin-1 in CGNs from WT and p75^DHEA^ mice in the presence or absence of NGF, as indicated. Scale bar, 10μm. Values were shown as mean±SD PLA puncta per cell from 3 independent experiments (30 neurons each). Unpaired t test with Welch’s correction.

Interestingly, internalization of p75^DHEA^ was unaffected in HCNs (Figure 8F), but significantly accelerated in CGNs (Figure 8G), a result that is in stark contraposition to the effects of the KKEA mutant. In agreement with this, p75^DHEA^ interacted more strongly with AP2B1 and less so with caveolin-1 than the wild type receptor in CGNs, as assessed by the PLA assay (Figure 8H and I). This bias is also in line with the enhanced activation of the RhoA pathway in the mutant neurons. We conclude that the RhoA and NF-kB pathways play fundamental but contrasting roles in the speed and extent of the internalization of the p75^NTR^ receptor in neuronal subpopulations of the mammalian brain.

## Discussion

Endocytosis and signaling are intertwined molecular processes (Sorkin and Zastrow, 2009). Much has been learnt during the past two decades about the many ways in which internalization and intracellular trafficking of plasma membrane receptors affect the quality and duration of their activity and downstream signaling (Hoeller et al., 2005; Sorkin and Zastrow, 2009; Roy and Wrana, 2005). Comparatively, less is know about the ways in which the signaling of plasma membrane receptors affects their own endocytosis and intracellular trafficking. Moreover, although p75^NTR^ internalization has been studied in some detail in peripheral neurons, our understanding of p75^NTR^ internalization in neurons of the mammalian brain has remained far behind. In this study, we uncover the unexpected differential internalization behavior of p75^NTR^ in HCNs and CGNs, and the importance of p75^NTR^ coupling to the RhoA and NF-kB pathways for the its internalization in these neuron subtypes.

We have identified a fundamental role of RhoA signaling in the internalization of p75^NTR^ in both HCNs and CGNs. Pharmacological or genetic inhibition of this pathway has been shown to affect, both positively and negatively, endocytosis of some plasma membrane proteins, such as the voltage-gated potassium channel Kv1.2 (Stirling et al., 2009) and the epidermal growth factor receptor (EGFR) (Kaneko et al., 2005), as well as synaptic vesicle recycling from the plasma membrane (Taoufiq et al., 2013). Mechanistically, these effects are thought to be mediated through phosphorylation of endocytic adaptor proteins (such as endophilin)by ROCK, a major downstream effector of RhoA signaling, and remodeling of the actin cytoskeleton through ROCK-mediated activation of the LIM/coffilin cascade (Kaneko et al., 2005; Guan et al., 2023). In these examples, however, RhoA/ROCK signaling affects the endocytosis machinery more generally, and the stimulus activating the pathway is either unknown or emanating from a secondary signal. On the other hand, examples of a receptor activity directly affecting its own endocytosis are scarce (Sorkin and Zastrow, 2009). Ligand stimulation of the EGFR or the NGF receptor TrkA has been shown to increase the pool of cellular clathrin that is associated with the plasma membrane in a Src kinase-dependent fashion, thus indirectly aiding EGFR and TrkA internalization (Wilde et al., 1999; Beattie et al., 2000). In the present study, we have shown that pharmacological inhibition of either RhoA or its effector ROCK reduced p75^NTR^ internalization. More importantly, however, genetically uncoupling the receptor from RhoA signaling resulted in both slower and overall reduced p75^NTR^ internalization as well as insensitivity to ligand-stimulated p75^NTR^ endocytosis, demonstrating that it is the connexion between the receptor and RhoA signaling that is specifically critical for its internalization, not simply the overall levels of RhoA activity in the cell. To our knowledge, this is a unique example of feed-forward regulation of receptor internalization through activation of RhoA signaling by the same receptor.

Our present results confirm the relatively fast internalization kinetics of p75^NTR^ in HCNs, as initially reported in our earlier study (Yi et al., 2021). Unexpectedly, a very different internalization behavior was found in CGNs, with a rate 5-fold slower, resembling the kinetics observed in peripheral neurons. This difference could not be attributed to differential cholesterol content in the membranes of the two neuron types as these were found to be very similar. In both HCNs and CGNs, p75^NTR^ internalization was enhanced by NGF stimulation to a similar extent, and was equally dependent on dynamin. On the other hand, a fundamental difference between the two neuron types was found in the engagement of p75^NTR^ with components of the clathrin and caveolin endocytosis machinery. Compared to HCNs, p75^NTR^ interaction with AP2B1 was weaker in CGNs, while interaction with caveolin-1 was stronger. As caveolin-mediated endocytosis is generally slower than clathrin-mediated (Mazumdar et al., 2021), this finding could explain the different kinetics of p75^NTR^ internalization in the two neuron types. The molecular bases of the differential engagement of the receptor with clathrin and caveolin components in HCNs and CGNs remain to be determined.

We used neurons expressing a mutant p75^NTR^ deficient in coupling to NF-kB signaling (p75^DHEA^)to assess the potential role of this pathway in p75^NTR^ internalization. We saw a dramatic enhancement in the speed of p75^NTR^ internalization in CGNs but no effect in HCNs, suggesting that RIP2 binding negatively regulates endocytosis of p75^NTR^ in CGNs. The lack of effect of the DHEA mutation in HCNs can be attributed to poor or negligible p75^NTR^ coupling to the RIP2/NF-kB signaling pathway. As we demonstrated in a previous study (Vicario et al., 2015), NGF treatment of HCNs fails to recruit RP2 to p75^NTR^ (in part due to very low levels of RIP2 expression in these neurons), nor does it induce IkB degradation or nuclear translocation of the p65^NF-kB^ subunit. It is interesting to note that JSH-23, an inhibitor of the nuclear translocation of the p65^NF-kB^ subunit, had no effect in the endocytosis of the receptor in either neuron subtype. Notably, despite its efficient blockade of NF-kB-mediated gene transcription, JSH-23 does not affect any of the upstream steps in the pathway (Shin et al., 2004). Although our results do not rule out a possible role of canonical components of the NF-kB signaling pathway in regulating p75^NTR^ internalization, including TAK1 and the IKK complex (IKKα and IKKβ), the fact that p75^DHEA^ showed stronger binding to RhoGDI and elevated RhoA•GTP levels suggest that this pathway may be responsible for the enhanced internalisation of this mutant receptor in CGNs. We have shown coupling to the RhoA pathway is critical for rapid and efficient p75^NTR^ internalization and, conversely, treatment with PMA, which increases RhoA activity, enhanced the endocytosis of this receptor. We thus favor the notion that the DHEA mutation accelerates p75^NTR^ internalization by preventing RIP2 binding, thereby allowing increased engagement with RhoGDI and RhoA activation.

In conclusion, this study demonstrates that p75^NTR^ internalizes with very different kinetics in two distinct neuronal subpopulations of the mammalian brain, reveals the importance of RhoA signaling in the endocytosis of this receptor, and demonstrates how a tug-of-war between RhoGDI and RIP2 for receptor engagement determine the cell type-specific p75^NTR^ internalization behaviors. The knock-in mouse strains expressing p75^KKEA^ and p75^DHEA^ mutant receptors described here will be useful in deciphering the contribution of these signaling pathways to the various functions of p75^NTR^ in mammalian physiology and neurodegeneration.

## Methods

### Mice

Animal care and experimental procedures were approved by Laboratory Animal Welfare and Ethics Committee of Chinese Institute for Brain Research (CIBR-IACUC-028). Mice were housed in a 12-h light–dark cycle and fed a standard chow diet. All mice (wild type and mutant)used for primary culture studies were male of C57BL/6 background. Constitutive knock-in p75^DHEA^ mice were generated by Taconic-Artemis, Germany; constitutive knock-in p75^KKEA^ were generated at the Chinese Institute for Brain Research.

### Cell line culture and transient transfection

NIH3T3 and HEK293T cells were cultured in Dulbecco’s Modified Eagle Medium (DMEM) (Gibco) plus 10% fetal bovine serum (Life Technologies)supplemented with penicillin-streptomycin (Life Technologies). The cells were transfected with the polyethylenimine (PEI) method. Briefly, cells were plated in a 10 cm tissue culture dish at a confluency of 2x 10^6^cells/ 10cm dish in normal growth media with antibiotics. Twenty-four hours after plating, transfection mix was prepared by mixing 10 µg of plasmid with 20 µg of PEI (1mg/ml) in DMEM. The transfection mix was left to stand at room temperature followed by addition dropwise into culture plates. 24 hours post-transfection, the cells were used for experiments. Where indicated, cells were treated with proNGF (Alomone) at 2.5ng/ml.

### Primary culture of hippocampal and cerebellar granular neurons

Primary hippocampal neurons (HCNs) were isolated from embryonic day 17.5 mouse embryos. Cerebellar granular neurons (CGNs) were isolated from postnatal day 3 mouse pups. Hippocampal structures or cerebellums were digested in Papain (Washington) for 30 min at 37°C and rinsed in Leibovitz’s-15 media. Neurons were triturated into a single cell suspension, counted with a hemocytometer, and then transferred to coverslips coated with 0.01% poly-D-lysine (Sigma-Aldrich) and 1 μg/ml mouse Laminin (R&D Systems). Cultured HCNs were maintained in serum-free defined Neurobasal media supplemented with B27 (Invitrogen) and GlutaMAX (Invitrogen) at 37°C in 5% CO2. CGNs were maintained in serum-free defined Neurobasal media supplemented with B27 (Invitrogen), GlutaMAX (Invitrogen) and KCl (Sigma-Aldrich) at 37°C in 5% CO2.

### Lentivirus production and transduction of CGNs

Lentiviruses expressing p75^NTR^ wild type, K303A, K346A, E349A, K303A/K346A, K303A/E349A and KKEA (K303A/K346A/E349A) utants were produced by transfecting HEK293T cells with pHAGE-IRES-eGFP constructs together with standard packaging vectors (pCMV-dR8.74 and pCMV-VSVG) by PEI based transfection method followed by ultra-centrifugation-based concentration of viral particles. Virus titer (T)was calculated based on the infection efficiency for HEK293T cells, where T = (P*N)/(V), T = titer (TU/µL), P = % of infection positive cells according to the fluorescence marker, N = number of cells at the time of transduction, V = total volume of virus used. Note TU stands for transduction unit. One day before transduction, CGNs were plated on PLL treated 24 well cell culture plate. On the next day, the neurobasal medium was aspirated and CGNs were incubated with 200µL of viral particles containing neurobasal media overnight. Next day the medium was removed and replaced with fresh neurobasal media and the cells were incubated for 1-2 days before experiments. Where indicated, cells were treated with proNGF (Alomone)at 2.5ng/ml.

### Antibodies

Primary antibodies used for immunofluorescence (IF)and immunoblotting (WB)were as follows:

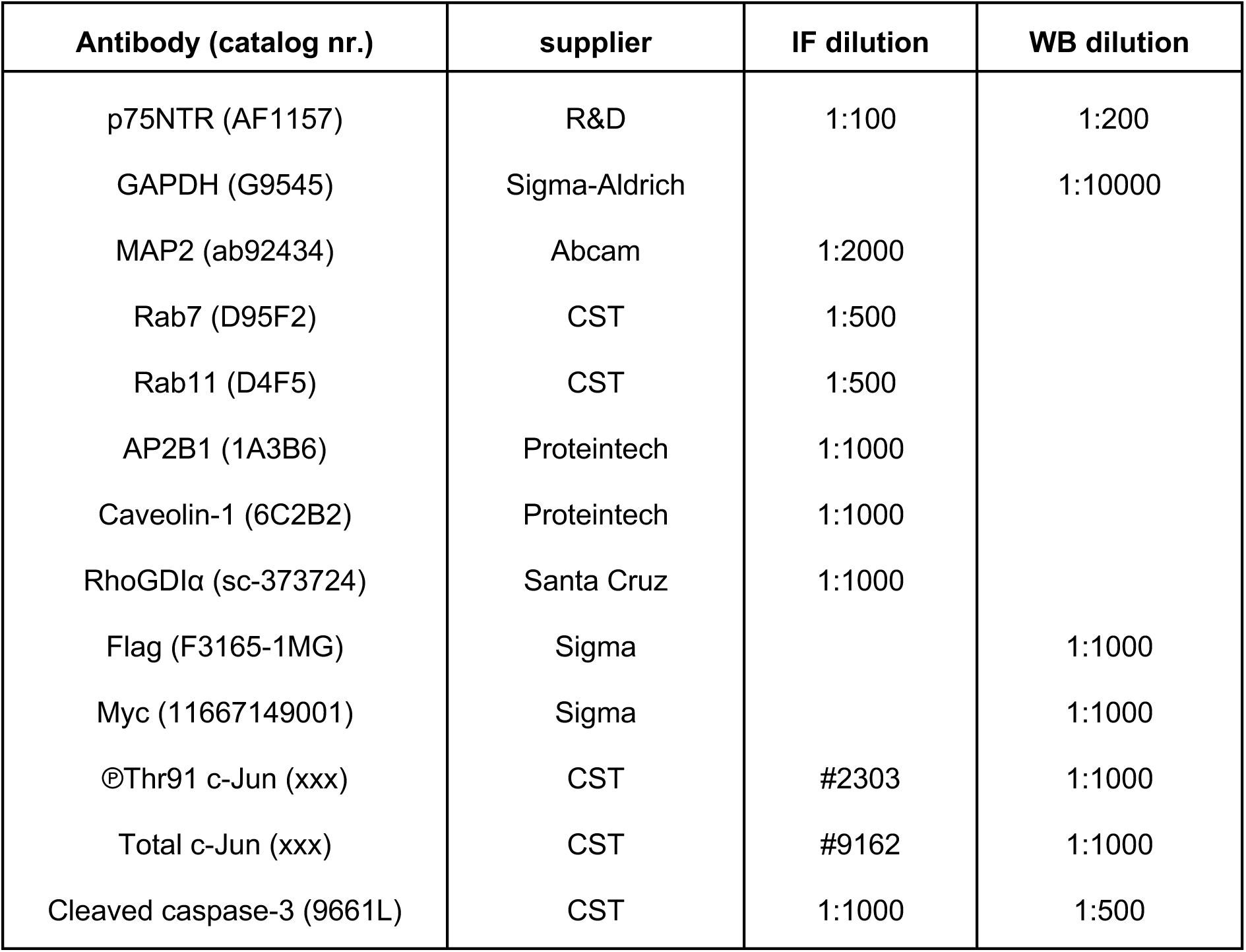

Secondary antibodies used for immunofluorescence were all four Invitrogen as follows: Donkey anti-Rabbit IgG Adsorbed Alexa Fluor™ 488 (A-21206), Donkey anti-Goat IgG Alexa Fluor™ 555 (A-21432) and Donkey anti-Chicken IgY Alexa Fluor™ 647 (A78952). All were used at a 1:1000 dilution.

### Pharmacology

Fasudil was used at 15μg/ml for 48h. Rhosin was used at 10μM for 48h. JNK-IN-8 was used at 3μM for 48h. Q-VD-OpH was used at 5μM for 48h. JSH23 was used at 10μM for 2h. Dynasore was used at 80μM for 30min. Gö6976 was used at 1μM for 30min. PMA was used at 2μM for 1h.

### Internalization assay

Primary antibody incubation was performed using goat anti-p75^NTR^ antibody (AF1157, R&D) diluted (1:100) in artificial cerebrospinal fluid (ACSF: 124 mM NaCl, 3.7 mM KCl, 1.0 mM MgSO4, 2.5 mM CaCl2, 1.2 mM KH2PO4, 24.6 mM NaHCO3, and 10mM D-glucose)for 1 h at 4°C to label surface p75. The cultures were then washed in ACSF and incubated at 37°C to allow internalization. NGF (R&D) was used at 50ng/mL together with primary antibody and 37°C incubation. At different time points, the internalization was stopped by quick wash in 0.2M formic acid (acid wash). Total staining (100%) was determined by direct fixation after antibody feeding, without acid wash. Baseline (t = 0 min) was obtained by acid wash directly after antibody labeling without 37°C incubation. The cultures were then fixed by addition of 4% PFA in PBS for 10 min at room temperature, washed with PBS, and incubated in blocking buffer for 30 min at room temperature. Labeled p75^NTR^ were detected by incubation with appropriate secondary antibodies at 1:1000 dilution in blocking buffer for 1h at room temperature. Cells were washed three times with PBS and mounted in Flouromount (Dako). In all experiments, cells were visualized on a Zeiss LSM900 confocal microscope.

### Proximity ligation assay (PLA)

Primary neurons were fixed for 10 min in 4% PFA, permeabilized, and blocked in 5% normal donkey serum and 0.2% Triton X-100 in PBS. To assess the effect of NGF (R&D), cells were treated with NGF at 50ng/mL for 1h. Cells were then incubated overnight at 4°C with primary antibodies in blocking buffer. The Duolink In Situ Proximity Ligation kit (Sigma) was used as per the manufacturer’s instructions with fluorophore-conjugated secondary antibody to recognize MAP2 included during the amplification step. The cultures were imaged with a Zeiss LSM900 confocal microscope to detect PLA signals. PLA puncta were quantified using ImageJ software.

### Co-immunoprecipitation and Western blotting

Cells were washed 3 times with sterile ice-cold PBS then lysed in lysis buffer (50 mM Tris/HCl pH 7.5, 1 mM EDTA, 270 mM Sucrose, 1% (v/v) Triton X-100, B mercaptoethanol and 60mM Octyl β-glucoside) containing protease and phosphatase inhibitors. Lysates were cleared at 10,000 × g pellet for 1 min at 4 °C. The protein concentration contained in the samples was determined using a BCA kit (Solarbio). For immunoprecipitation, 1μg of p75^NTR^ antibody was added to a total of 500 μg protein (1 mg/mL in lysis buffer) and was incubated overnight on an orbital shaker at 4 °C. On the next day, 25 μL bed-volume of ethanolamine treated protein G sepharose resin (Pierce) was added to the lysate, and mixtures were incubated for 2 hours at 4 °C with rotation. The resin was washed once with wash buffer 1 (50mM Tris, 250mM Sucrose, 5mM MgCl2, 0.15M NaCl, 2% Igepal, 200mM ethanolamine, pH 10.5) and washed 3× with the wash buffer 2 (20mM Tris, pH 7.5, 250mM Sucrose, 5mM MgCl2, 0.15M NaCl, 2% Igepal), then boiled in 30μL 2× SDS sample buffer, and eluted proteins were analyzed by SDS/PAGE followed by Western blotting. 20μg of clarified total cellular lysate was loaded per well to be separated on an SDS-PAGE followed by Western blotting on polyvinylidene difluoride (PVDF)membranes. Western blots were probed using HRP conjugated secondary antibodies, and detected using ECL detection kit (Cell Signaling).

### RhoA and NF-kB activation assays

On the day of harvest, cells were rinsed immediately in ice-cold phosphate-buffered saline (PBS, Sigma)prior to lysate collection. Cells were stored in G-LISA lysis buffer with manufacturer’s protease inhibitors (Cytoskeleton). The absorbance at 490 nm measures the active GTP-bound RhoA in the neuronal lysates that have bound to the Rhotekin RhoA binding domain immobilized to the 96 well plate. Protein concentrations were determined using Precision Red Advanced Protein Assay Reagent with absorbance at 600 nm measured spectrophotometer. Lysates were incubated in wells for 30 min at 4 °C with shaking, then rinsed and sequentially incubated with primary antibodies to RhoA followed by HRP secondary antibodies (Cytoskeleton), and the colorimetric reaction was performed for 15 min at 37 °C, which was measured using a microplate plate reader (BioTek). To assess NF-kB activation, we used an assay (Active Motif products, #43296)in which a colorimetric readout measures activated NF-kB p65 subunit on whole cell extracts. Whole cell lysates from cultures of CGNs were applied to a 96-well plate to which oligonucleotide containing a p65 NFκB consensus DNA binding site has been immobilized. The colorimetric readout at 450 nm measures activated NFkB p65 on whole cell extracts bound to the oligonucleotide immobilized to the 96 well plate.

### Immunohistochemistry

HCNs and CGNs cultured on coverslips were briefly washed in PBS, fixed for 10min in 4% paraformaldehyde solution, and blocked in PBS containing 5% donkey serum and 0.2% Triton X-100. Fixed cells were then incubated overnight at 4°C with the appropriate antibodies, followed by incubation in fluorophore-conjugated secondary antibodies. Coverslips were mounted onto microscopy slides using Flouromount (Dako). Confocal laser scanning microscopy was performed on a Zeiss LSM900 microscope.

### Assessment of cellular cholesterol

Cholesterol content of plasma membrane was detected by filipin staining. Primary neurons were fixed for 10 min in 4% PFA. Cells were stained with Filipin Ⅲ solution (1:40, Sigma-Aldrich, SAE0087) with 10% normal donkey serum for 1h. To permeabilize the cells, 0.2% Triton X-100 was applied together with the Filipin staining. TO-PRO3 was used at a dilution of 1:1000 for 15 minutes at room temperature. Cholesterol level of cell extraction was detected by AmplexRed cholesterol kit (Invitrogen, A12216). Lipid extraction was following the Bligh and Dyer method with modifications as follows: the cultured cells were homogenized with a mixture of chloroform, methanol, and water (2:1:0.8 v/v/v). Phase separation was induced by adding equal volumes of chloroform and water to the homogenized mixture, followed by centrifugation at 1000rpm, 5min at room temperature. The lower phase containing the lipids was used for cholesterol detection following the instructions of AmplexRed cholesterol kit.

### Image analysis

Image processing and quantification were performed using ImageJ software, with colocalization analysis conducted through the JACoP plugin for FIJI. In the internalization assay, at least five fields were captured and quantified for each replicate coverslip. The mean fluorescence intensity of p75^NTR^ staining within the MAP2-positive area was quantified. All data were normalized to the total staining group. For PLA image analysis, the number of PLA puncta per cell was counted using ImageJ, with a constant threshold applied across all images. Microscope parameters were maintained consistently for all images to allow direct comparison.

### Statistical analysis

GraphPad Prism (versions 4 or 8; GraphPad Software Inc, San Diego, CA, USA)were used for statistical analyses. Experimental data was collected from multiple experiments and reported as the mean ± SD or mean ± SEM described in the corresponding Figure Legends. Statistical significance was calculated using the one-way ANOVA followed by Tukey’s multiple comparisons test for all data except those shown in Figures 2, 5A, and 7 which used the Student t-test. P value of less than 0.05 was considered statistically significant.

## Acknowledgements

The authors would like to thank Lei Wang, Jocelyn Jia, Yankui Fu and Shuo Zhang for technical and admin assistance. This work was supported by research grants to C.F.I. from Peking University, Chinese Institute for Brain Research, Beijing, and Swedish Research Council (Vetenskapsrådet, contract nr. 2024-03222); and a startup grant to M.X. from Swedish Research Council (Vetenskapsrådet, contract nr. 2021-01805).

## Author contributions

X.L. performed all experimental work, except biochemistry assays shown in Figure 4 (performed by A.R.)and PLA assays shown in Figure 5 (performed by Z.F.). X.L., Z.F and C.F.I. analyzed the data; X.L. prepared the figures; M.X. co-directed the project and corrected the manuscript; C.F.I. conceived the project, directed the research and wrote the manuscript.

## Supplementary figure legends

**Supplementary Figure S1.**
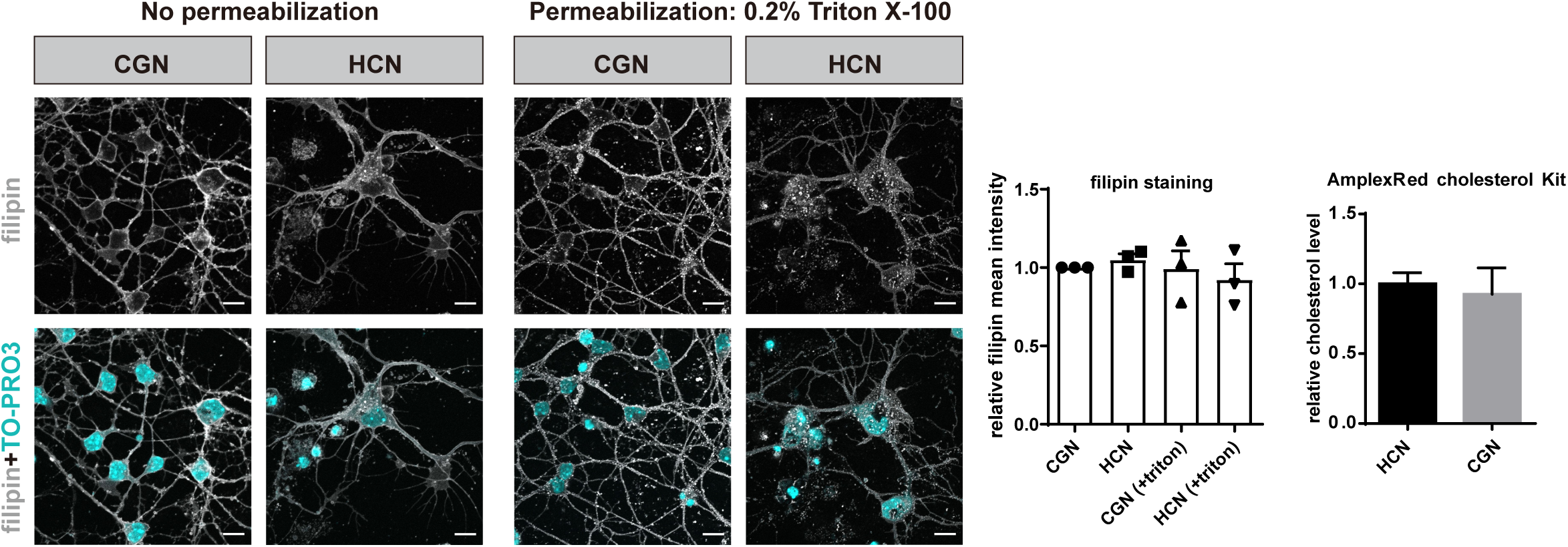
Similar cholesterol content in HCNs and CGNs. (A) Visualization of cholesterol in HCNs and CGNs (as revealed by Filipin staining with counterstaining of neuronal nuclei with TOPRO3. (B) Quantification of Filipin staining in cultured HCNs and CGNs before and after permeabilization with Trion X100. Results are presented as mean ± SEM of three independent experiments, each performed in duplicate, normalized to the CGN group. Results were not significantly different as judged by unpaired t test with Welch’s correction. (C) Quantification of cholesterol levels in HCNs and CGNs using AmplexRed cholesterol detection kit. Results are presented as mean ± SEM of three independent experiments, each performed in duplicate. Results were not significantly different as judged by unpaired t test with Welch’s correction.

**Supplementary Figure S2.**
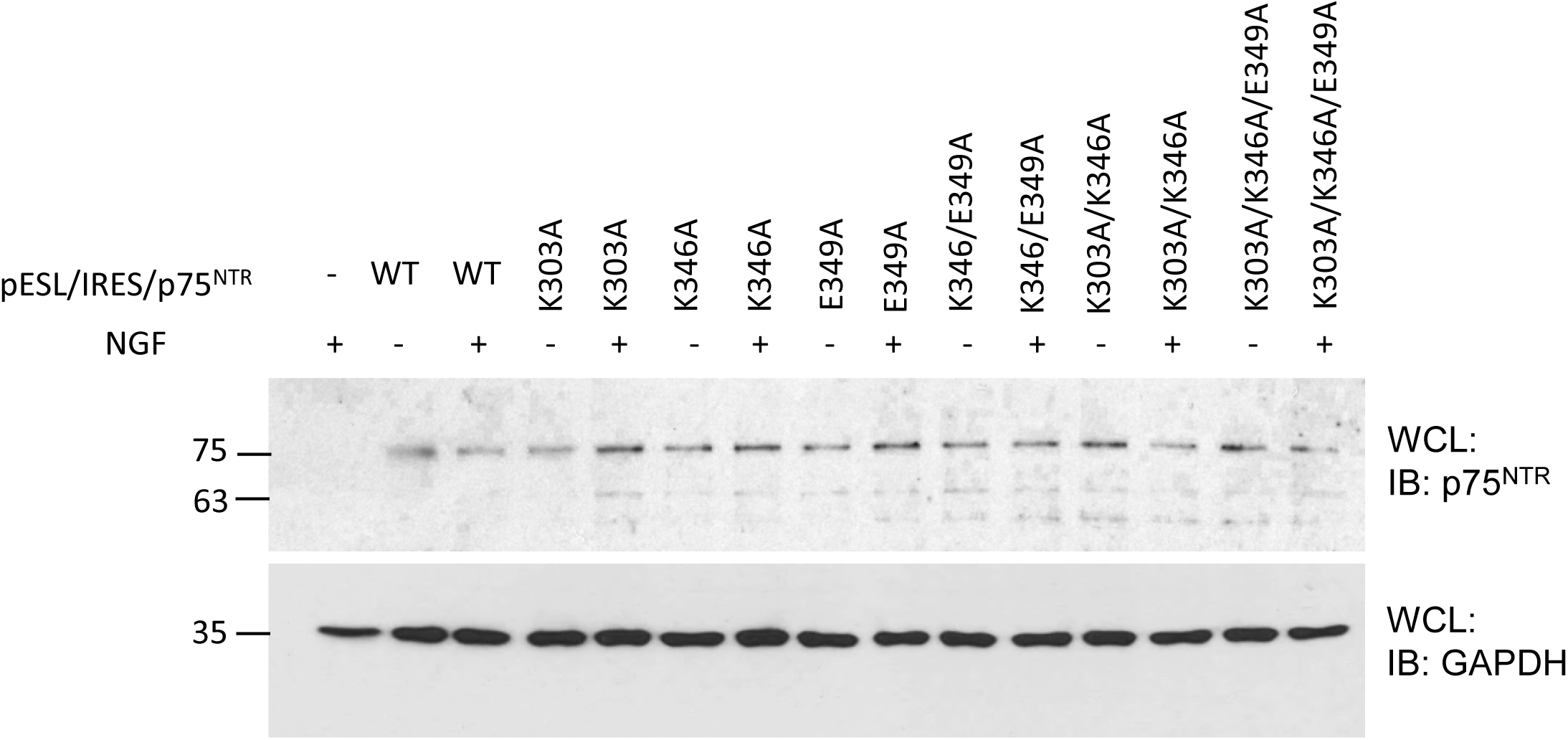
Comparable levels of mutant p75^NTR^ expression in lentiviral-transduced CGNs. Western blotting of extracts from CGNs lacking p75^NTR^ (knock-out)that were reconstituted with the indicated constructs via lentivirus transduction, probed for p75^NTR^ and reprobed for GAPDH as loading control.

**Supplementary Figure S3.**
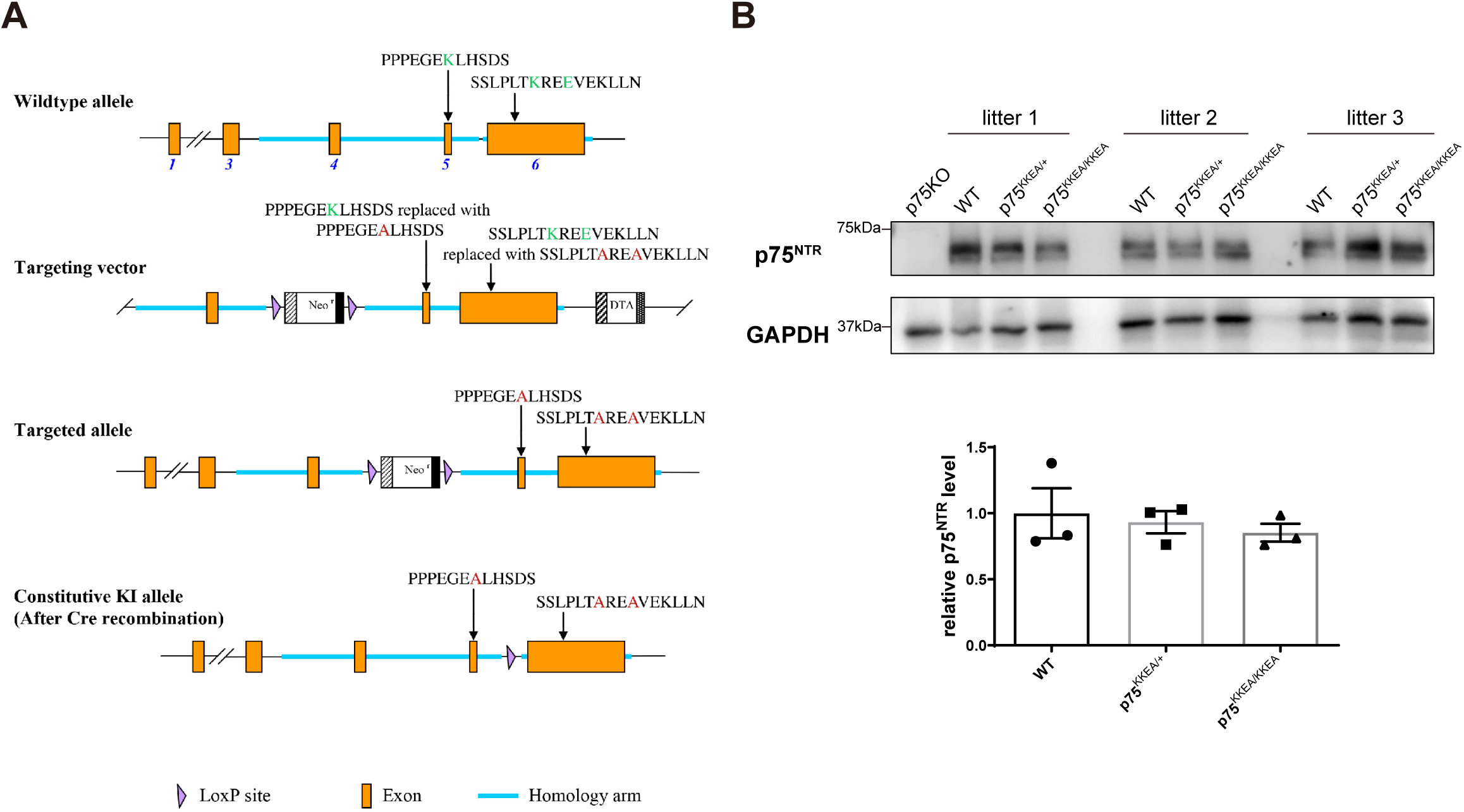
Generation of knock-in KKEA p75^NTR^ mice. (A) Schematic of the *p75ntr* locus (not at scale)with strategy for generation of the KKEA mutant knock-in allele. (B) p75^KKEA^ mutant mice express normal levels of the mutant receptor in postnatal and adult brain. Shown are immunoblots of p75^NTR^ of brain extract from P7 pups of different genotypes. p75^NTR^ was detected by AF1157 antibody. The lower panel shows GAPDH as a control for equal loading. Quantification shows mean±SEM. N=3. Unpaired t test with Welch’s correction.

**Supplementary Figure S4.**
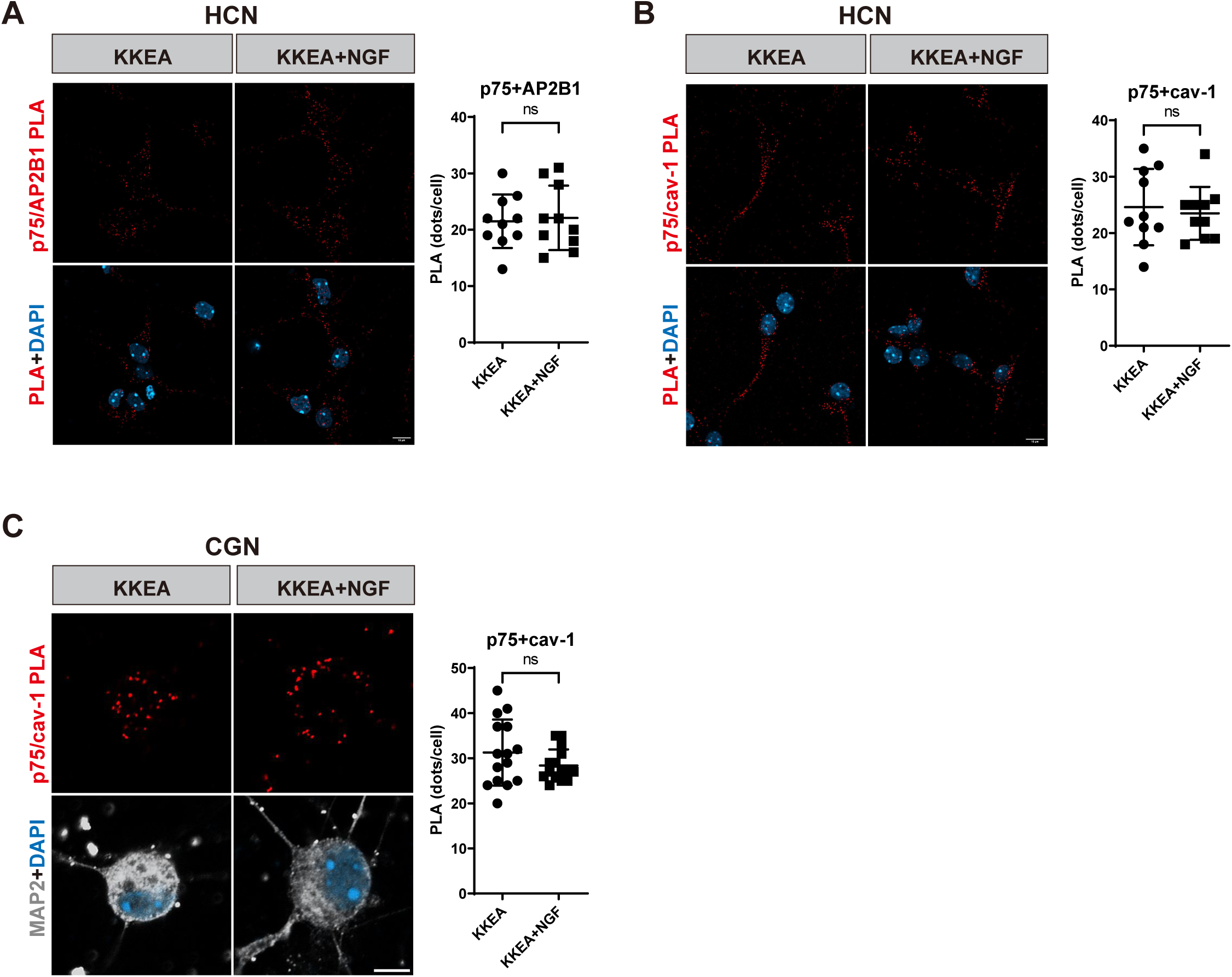
Interaction of p75^KKEA^ with components of the clathrin and caveolin endocytosis pathways is not affected by NGF treatment. (A) PLA assay between p75^NTR^ and AP2B1 in KKEA HCNs in the presence or absence of 50ng/mL NGF. Scale bar, 10μm. Values were shown as mean±SD PLA puncta per cell from 10 neurons from duplicate coverslips. Unpaired t test with Welch’s correction. ns, not significant. (B) PLA assay between p75^NTR^ and caveolin-1 in KKEA HCNs in the presence or absence of 50ng/mL NGF. Scale bar, 10μm. Values were shown as mean±SD PLA puncta per cell from 10 neurons from duplicate coverslips. Unpaired t test with Welch’s correction. ns, not significant. (C) PLA assay between p75^NTR^ and caveolin-1 in KKEA CGNs in the presence or absence of 50ng/mL NGF. Scale bar, 5μm. Values were shown as mean±SD PLA puncta per cell from 15 neurons from duplicate coverslips. Unpaired t test with Welch’s correction. ns, not significant.

**Supplementary Figure S5.**
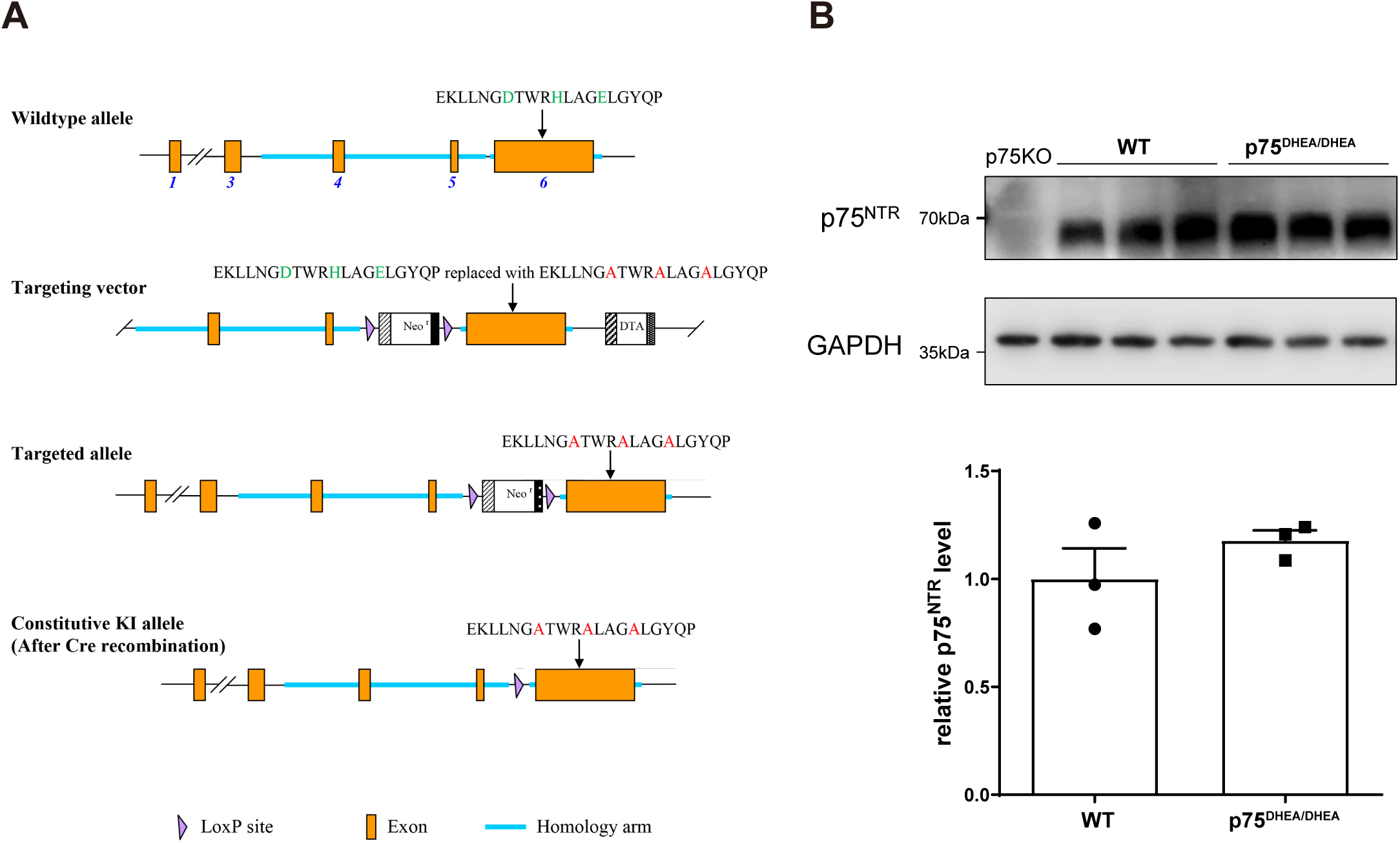
Generation of knock-in DHEA p75^NTR^ mice. (A) Schematic of the *p75ntr* locus (not at scale)with strategy for generation of the KKEA mutant knock-in allele. (B) p75^DHEA^ mutant mice express normal levels of the mutant receptor in postnatal and adult brain. Shown are immunoblots of p75^NTR^ of brain extract from E17.5 embryos of different genotypes. p75NTR was detected by AF1157 antibody. The lower panel shows GAPDH as a control for equal loading amount. Quantification shows mean±SEM. N=3. Unpaired t test with Welch’s correction.

